# Microbial alginate foraging is conserved in geographically and taxonomically distinct ruminant microbiomes

**DOI:** 10.1101/2024.12.17.628917

**Authors:** Alessandra Ferrillo, Jeffrey P. Tingley, Marissa L. King, Alemayehu Kidane, Barinder Bajwa, Xiaohui Xing, Tina Johannessen, Alexsander Lysberg, Liv Torunn Mydland, Margareth Øverland, Greta Reintjes, Anna Y. Shearer, Leeann Klassen, Kristin E. Low, Trushar R. Patel, Stephanie A. Terry, Phillip B. Pope, D. Wade Abbott, Live H. Hagen

## Abstract

Seaweed plays a crucial role in carbon cycling and is expected to be a valuable resource for sustainable biomass, with applications in biofuel production, human nutrition, and animal feed. Although seaweed has historically been used as a feed source for livestock grazing near coastlines, the process by which it is digested in the rumen remains unknown. Here, we show how the brown algae *Saccharina latissima* is catabolized in the rumen ecosystem of two different species using *in vivo* and *in vitro* experimental systems. We determined that the ruminal decomposition of alginate, a prominent component of the brown algae cell wall, requires microbial catabolic pathways complete with alginate lyases and transport proteins. Evidence of digestion was obtained through a combination of animal models, bacterial imaging, multilayered meta-omics, and enzyme biochemistry. The evolution of and implications for acquisition of ‘alginate utilization loci’ within geographically and taxonomically distinct ruminants are considered.

**Graphical abstract:** *Saccharina latissima* is a brown alga commonly found in the North Atlantic, Arctic and Pacific oceans. *S. latissima* was collected from the west coast and Canada and Norway for microbiome studies. Alginate constitutes a substantial portion of the cell wall of *S. latissima* (SL), and its digestion requires a specific set of enzymes, alginate lyases. We investigated if and how *S. latissima* is metabolized in geographically distinct rumen ecosystems through *in vivo* lamb feeding experiments (2.5 and 5% inclusion, DM basis) and *in vitro* cattle-based rumen simulation technique, RUSITEC, experiments (up to 50% inclusion). Evidence supporting ruminal degradation of alginate was explored using a combination of multilayered meta-omics, physiology (fluorescently labelled *S. latissima* hot water extracts (FLA-SLAT)) and biochemical characterization of PL6 alginate lyases.

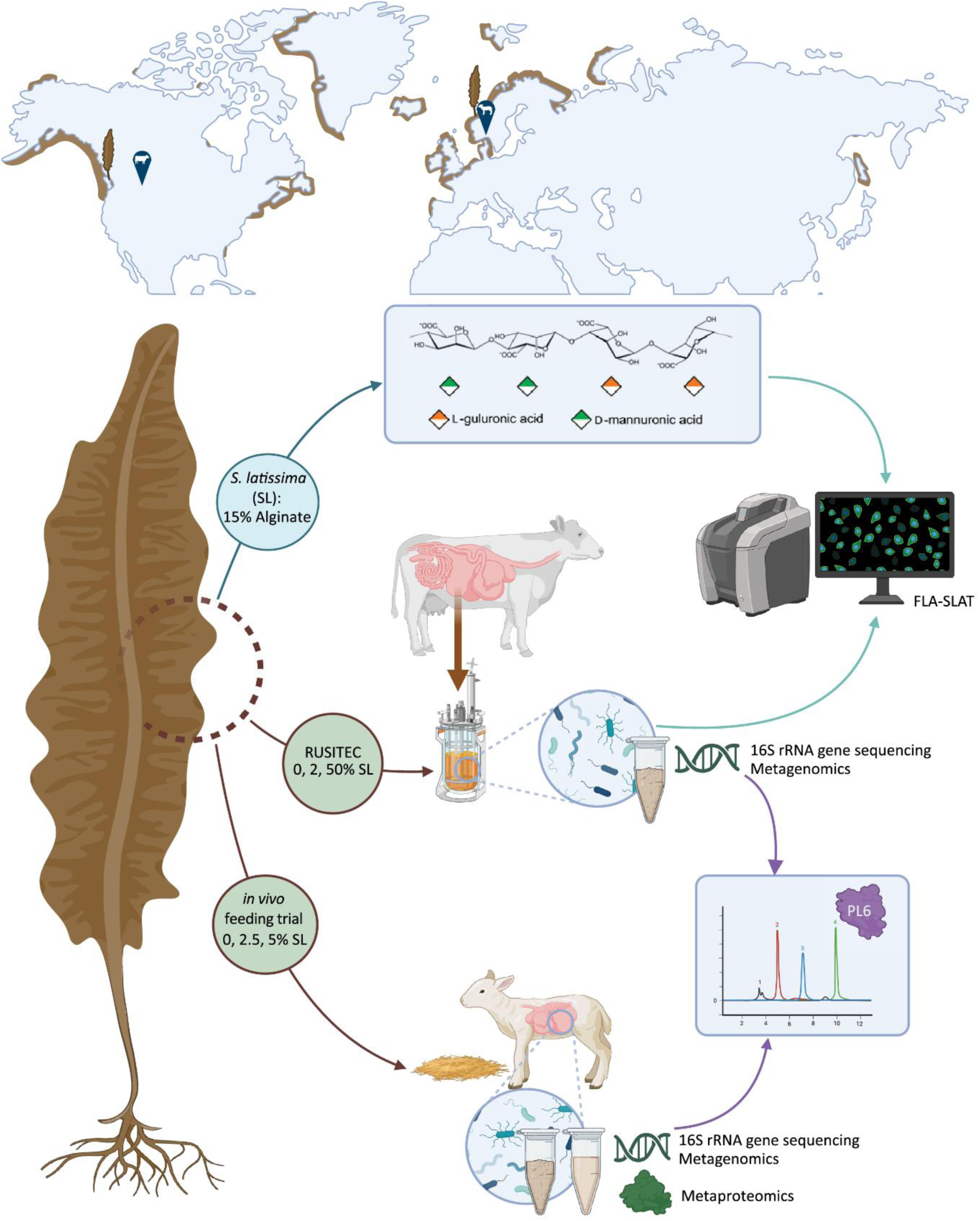

## INTRODUCTION

The effects of climate change, such as extreme drought and flooding, are deleterious for food production and reduce access to cultivatable land. Thus, seeking solutions from the ocean to address the challenges associated with the harvest of terrestrial food and feed is garnering interest. Cultivated and wild-harvested macroalgae are emerging as a promising source of biomass for future application in biofuel production, human nutrition, and animal feed ^1^. Certain varieties of marine macroalgae, such as kelp, grow faster than terrestrial feed crops ^2, 3^, and create highly productive coastal forests that sequester substantial amounts of carbon globally while improving habitat for other marine life ^4^. Importantly, macroalgae cultivation does not rely on freshwater, nor compete with arable land or land areas dedicated to conservation, which makes it an attractive source of sustainable biomass. Algal biomass is also rich in micronutrients and proteins, thus has potential to supplement livestock diets ^5^.

The brown macroalga *Saccharina latissima* (‘sugar kelp’) belongs to the *Laminariaceae* family and is widely distributed across temperate coastal waters, including the North Atlantic and North Pacific. Like most brown macroalgae, the cell wall of *S. latissima* is largely comprised of structural polysaccharides such as alginate, cellulose, and fucoidans, which together account for 30-50% of the algal total dry weight ^6^. Depolymerization of alginate is catalyzed by a highly specialized group of polysaccharide lyase (PL) enzymes, called alginate lyases, that cleave glycosidic bonds through β-elimination, resulting in unsaturated products ^7^. A number of alginate lyases have been studied from various organisms, primarily originating from marine environments, including marine algae ^8^, fungi ^9^, viruses ^10^, marine mollusks ^11, 12^ and human gut bacteria ^13^. Intriguingly, previous studies have highlighted possible transmission of seaweed-degrading enzymatic machinery from marine bacteria to the human gut microbiome ^14, 15, 16, 17^. This includes alginate lyases ^13^ and polysaccharide utilization loci (PULs) for red-algal galactan degradation in human gut bacteria ^15, 18^.

In recent years, the practice of adding seaweed to livestock feed has gained momentum due to the potential of certain species to reduce enteric methane emissions ^19, 20^. However, feeding seaweed to cattle is not a new strategy; farmers in coastal areas have historically collected kelp to supplement the livestock feed in years with poor harvests ^21^. In some instances, animals have survived almost entirely on seaweed ^22, 23^. While some ruminant populations, such as the North Ronaldsay sheep, have consumed seaweed for generations, the evolutionary events that endowed intestinal microorganisms with the potential to convert their chemically complex polysaccharides into a source of nutrition for their hosts, are as-yet unknown. In this study, we investigated the adaptation of two geographically distinct ruminant microbiomes to dietary kelp alginates. To evaluate microbiome responses, we first analyzed rumen samples from an *in vivo* feeding experiment, where naïve lambs were fed 2.5 and 5% *S. latissima* on dry matter (DM) basis. Through meta-omic analyses, we demonstrate the presence of functionally active rumen-associated *Prevotella* populations with the enzymatic machinery required to catabolize brown seaweed alginate through ‘alginate utilization loci’ (AULs). These results were validated in an unrelated rumen ecosystem using the rumen simulation technique (RUSITEC), where rumen content from cattle was incubated with *S. latissima* at low (2.0%) and high (50%) inclusion rates. Interactions between alginate and microbial cells were confirmed using fluorescently labelled polysaccharides (FLAPS) and extracted rumen communities. The activity of *Bacteroidota* AULs was further validated using recombinant enzymes derived from both lamb and cattle datasets. Alginate-consuming microbial populations were present across different production systems and different host species, yet with high homology between alginate lyases within the identified AULs. Our results underscore the remarkable ability of rumen microbial inventory to rapidly adapt to novel and complex polysaccharides.

## RESULTS

### Algal polysaccharides in *S. latissima*

A complete understanding of ruminal kelp digestion requires knowledge of the chemical and structural diversity of their polysaccharides. To determine the linkage composition, with a focus on assessing alginate content and the ratio of D-mannuronic (M) and L-guluronic (G) acids (**Figure 1A**), *S. latissima* harvested off west coast of Canada was subjected to GC-MS based glycosidic linkage analysis using a recently established procedure optimized for unfractionated cell wall of brown seaweeds ^24^. The identification of all permethylated alditol acetates (PMAAs), including those from 4-Gul*p*A and 4-Man*p*A, was based on their characteristic MS fragmentation patterns and retention times (**Figure 1B** and **C**). The relative abundance of linkages identified from *S. latissima* (**Supplementary Table S1**; **Supplementary Figure S1**) was used to estimate the relative compositions of total cell wall polysaccharides including cellulose, mixed linkage glucan (MLG), laminarin, fucoidan, and alginate (**Figure 1D**). Cellulose was found to be the most abundant polysaccharide, as indicated by the dominance of 4-Glc*p*, which is consistent with our comparative analysis of various brown seaweeds ^24^. Fucoidans, whose linkages were assigned based on previous study ^25^, exhibited a highly diverse linkage pattern, including all Fuc*p* (17.2%), all 2-Man*p* (4.8%), all 6-Gal*p* (3.8%), t-Gal*p* (trace), 3,4-Gal*p* (1.4%), 4-Glc*p*A (2.4%), 3-Glc*p*A (2.6%), t-Xyl*p* (1.5%) linkages, together representing 33.5% of the cell wall polysaccharides in *S. latissima*. Alginate constituted 14.9% of the total relative abundance, with Gul*p*A linkages (10.2%) being more prevalent than Man*p*A linkages (4.7%), suggesting an M/G ratio of 0.46. The laminarin composition was calculated to be 1.3%, based on the sum of 3-Glc*p* and 3,6-Glc*p*, multiplied by two to account for the corresponding t-Glc*p* of 3,6-Glc*p* side chains. Of note, MLG linkages may be confounded with those found in laminarin and cellulose.

**Figure 1:**
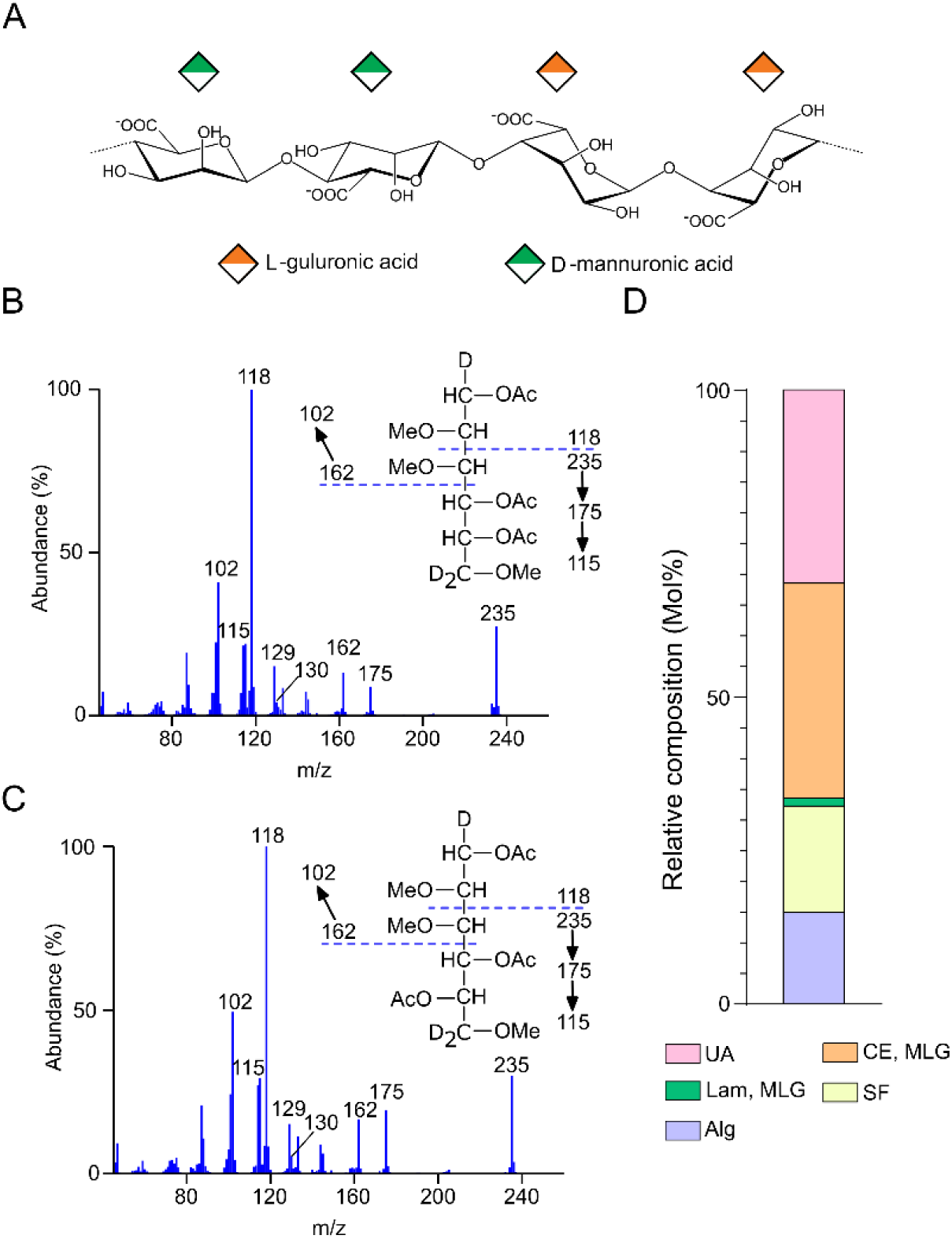
Linkage analysis and polysaccharide estimation of the brown algae *S. latissima* cell walls. A) Alginate is a high molecular weight, unbranched matrix polymer composed of two uronic acids: α-L-guluronic acid and β-D-mannuronic acid. These are linked by 1,4-glycosidic bonds and arranged in blocks of either homodimers (polyM or polyG) or heterodimers (polyMG). B) EI-MS spectra and ion fragmentation patterns for 4-Gul*p*A and C) 4-Man*p*A from PMAAs of *S. latissima* alginate. D) Estimated polysaccharide composition (Mol%) of cell walls extracted from *Saccharina latissima* (n=2). The following abbreviations were used for their respective polysaccharides: SF (fucoidans – All Fuc*p*, All 2-Man*p*, All 6-Gal*p*, t-Gal*p*, 3,4-Gal*p*, 4-Glc*p*A, 3-Glc*p*A, t-Xyl*p*) assigned according to the literature^25^; CE/MLG (cellulose, mixed linked glucan – 4-Glc*p* + 4,6-Glc*p* + (t-Glc*p* – 3,6-Glc*p*)); Alg (alginate – All Gul*p*A and Man*p*A linkages); Lam/MLG (laminarin, mixed linkage glucans – 3-Glc*p* + 3,6-Glc*p*\*2); UA (unassigned – all remaining linkages).

### Niche specialization dictates decomposition of brown algae polysaccharides in naïve ruminants

In our initial study of the rumen microbiome from Norwegian White lambs (n=24), 16S rRNA gene analysis was conducted on both fluid and fiber-attached rumen samples obtained from all animal dietary groups participating in the month-long feeding trial, including control, low seaweed (2.5% *S. latissima* on DM basis) and high seaweed (5.0% *S. latissima* on DM basis) (**Supplementary Table S2**). As expected, notable differences in the community structure between the fluid and fiber-attached samples were observed across all three dietary groups. However, no definitive evidence indicating a macroalgae-induced alteration in the microbiome structure was identified (**Supplementary Figure S2**). In the bovine RUSITEC experiment, where the range of seaweed inclusion was 2.0 and 50% *S. latissima* (DM basis), significant changes to rumen microbial communities were observed between time points (**Supplementary Figure S3A**; ANOSIM and PERMANOVA, *P* = 0.001) and 50% *S. latissima* inclusion (**Supplementary Figure S3B**; ANOSIM, *P* = 0.005; PERMANOVA, *P* = 0.008) compared to the control forage (**Supplementary Table S3**). RUSITEC communities supplemented with *S. latissima* also showed lower Shannan and inverse Simpson diversity values compared to control communities, although the Chao1 richness index was not significantly altered over time, or by seaweed supplementation (**Supplementary Figure S3C**). Notably, an evident increase in the relative abundance of *Bacteroide*s was seen at the 8-day time point in the 50% *S. latissima* treatment (**Supplementary Figure S3D**). In both rumen systems, digestibility of seaweed was demonstrated and there was no observed impact on other metabolic metrics (**Supplementary Text S1).**

Although the overall rumen community structure seemed unaffected by dietary supplementation of seaweed when given at biologically relevant doses, we wanted to further elucidate the genetic potential to utilize brown seaweed within the microbiomes. For this, shotgun metagenomic sequencing and MAG reconstruction were carried out on lamb rumen samples from all three dietary groups, as well as for the control and 50% *S. latissima-*treated RUSITEC communities. Reconstruction of the lamb rumen metagenome yielded 260 MAGs (named “MAGX_OV_”) of medium to high quality, defined by the MIMAG standard for completeness and contamination ^26^. The majority of these MAGs were affiliated with the *Bacteroidota* phylum (n=148), followed by *Bacillota* (n=36) and *Methanobacteriota* (n=7) (**Figure 2A**). When investigating the gene content of these MAGs, we identified enzymes annotated as alginate lyases encoded within 28 MAGs (**Extended Data 1**). Most of these MAGs were associated with members of *Bacteroidota*, along with *Fibrobacterota* and *Verrucomicrobiota*. Due to the structural composition of alginate, alginolytic systems often appear in gene clusters, or AULs, in Gram-negative bacteria; these systems consist of Sus-like proteins and alginate lyases with complementary specificities ^16, 27, 28^. We therefore extended our gene search to include the co-occurrence of TonB-dependent receptors (SusC) and SusD-like substrate-binding proteins, which are archetypical components of PULs ^29, 30, 31^. This refinement led to the identification of ten *Bacteroidota* MAGs, of which four were classified as *Prevotella* species, five assigned to the *Bacteroidales* RC9 clade, and one to the *Paludibacteriaceae* family (**Figure 2A**).

**Figure 2:**
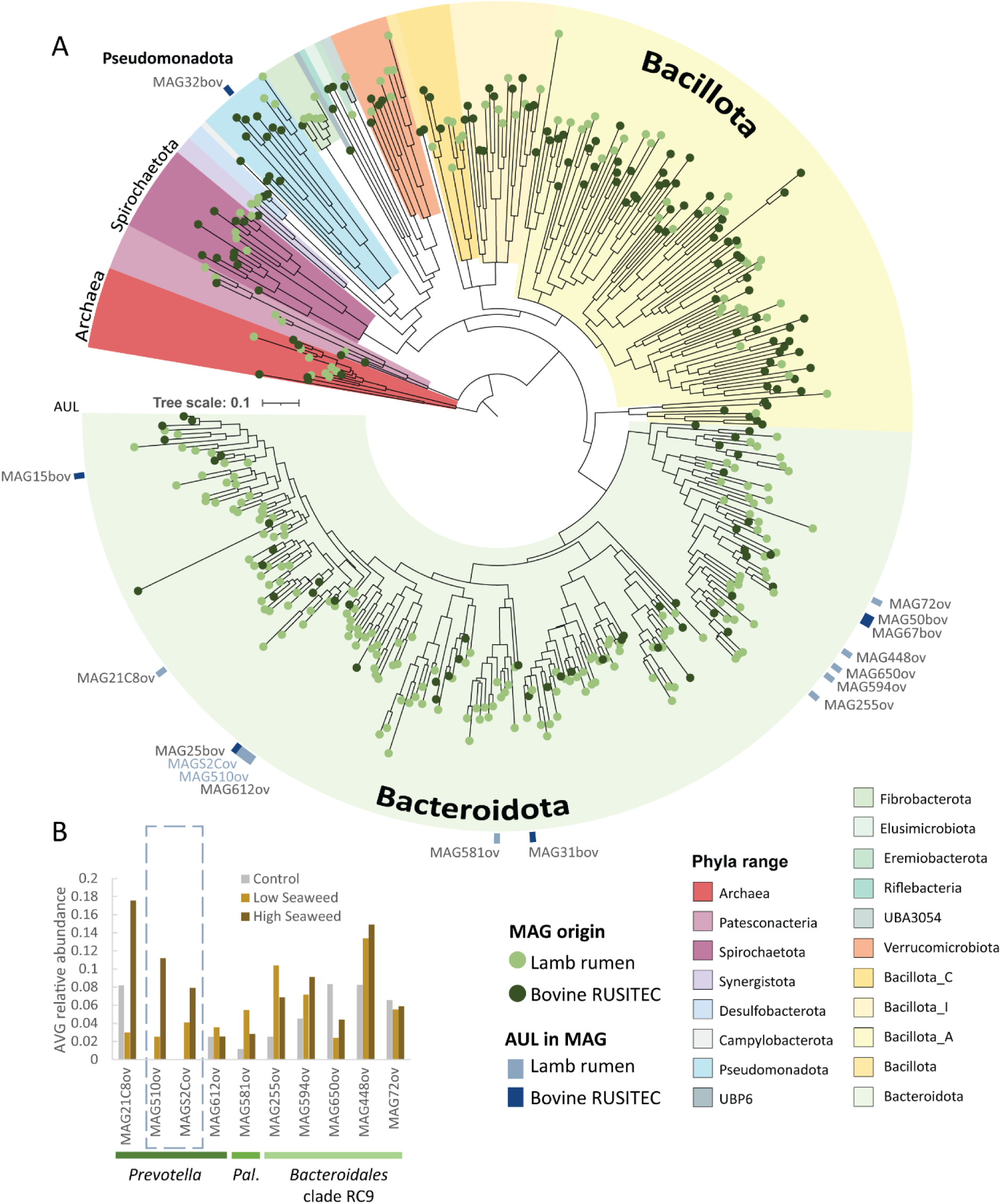
Rumen microbes encode enzymes for alginate utilization. A) Phylogenetic tree of MAGs recovered from lamb rumen (light green circle on branch) and the bovine RUSITEC (dark green circle on branch) microbiomes. In total 16 MAGs, of which all except one (MAG32_BOV_) are affiliated to the *Bacteroidota*-phylum, encoded AUL for alginate decomposition. B) Relative abundance of the lamb rumen MAGs encoding alginate lyases were calculated for each sample using CoverM and displayed as the average of each diet group. Two closely related *Prevotella* MAGs harboring AUL (MAGS2C_OV_ and MAG510_OV_) demonstrated dose-dependent increase in relative abundance in response to inclusion of *S. latissima* in the diet. Pal.; *Paludibacteraceae*.

The genomes recovered from the RUSITEC systems accounted for another 132 MAGS (named “MAGX_BOV_”), with a noticeable fraction being *Bacillota* (n=64) and *Bacteroidota* (n=32) (**Figure 2A**). Similar to the lamb microbiome, the majority of alginate lyases were found within *Bacteroidota* phylum members originating from the *S. latissima-*treated RUSITEC system (**Extended Data 1**); six *Bacteroidota* MAGs contained alginate lyases, five of which also harbored intact AULs (**Figure 2A**). Notably, an alginate utilization cluster (AUC) was also identified within MAG32_BOV_ annotated as *Brevimundis bullata* (87% complete; contamination <1%), which does not have the traditional SusC/D system.

Intriguingly, among the lamb-originating MAGs encoding alginate lyases, two closely related *Prevotella*-affiliated MAGs, MAG510_OV_ and MAGS2C_OV_, demonstrated a pronounced increase in relative abundance in response to increasing levels of *S. latissima* in the diet. In contrast to the other potential alginate-degrading genotypes, none of these two *Prevotella* MAGs were detected in fiber-attached rumen samples from the control diet group (**Figure 2B**), further indicating that their presence correlates with *S. latissima* inclusion in the diet.

### Functionally active alginate consuming *Prevotella* populations

To investigate whether the putative alginate-degrading populations were functionally active, we analyzed the metaproteome of rumen samples from all lambs in the control and high *S. latissima* (5%) dietary groups. After filtering, our metagenome-centered metaproteomic analysis resulted in detection of 10,312 protein groups that mapped back to our sample-specific database (**Extended Data 2**). Among these protein groups, we detected two groups annotated as alginate lyases (**Figure 3A**), suggesting a role in the depolymerization of alginate. Notably, these two alginate lyases, both PL6, mapped back to the two *Prevotella* MAGs (MAG510_OV_ and MAGS2C_OV_) that demonstrated dose-dependent increase in relative abundance in response to inclusion of *S. latissima* in the diet. While the peptides constituting one of the detected PL6 enzymes appeared to uniquely match MAGS2C_OV_, the other PL6 undistinguishably mapped back to protein sequences found in both *Prevotella*-affiliated MAGs. Alignment of the full-length protein sequences within the shared PL6 protein group revealed a high degree of sequence similarity (BLASTp: 96% id), suggesting a common ancestral origin and functional redundancy. The label-free quantification (LFQ) of the detected proteins confirmed that the two PL6 enzymes were enriched in the animal group fed with high inclusion of *S. latissima* (**Figure 3B**). Of note, in both MAG510_OV_ and MAGS2C_OV_, the genes of the expressed PL6 enzyme were neighboring another gene encoding an enzyme from a known alginate lyase family; PL17 (no protein detection).

**Figure 3:**
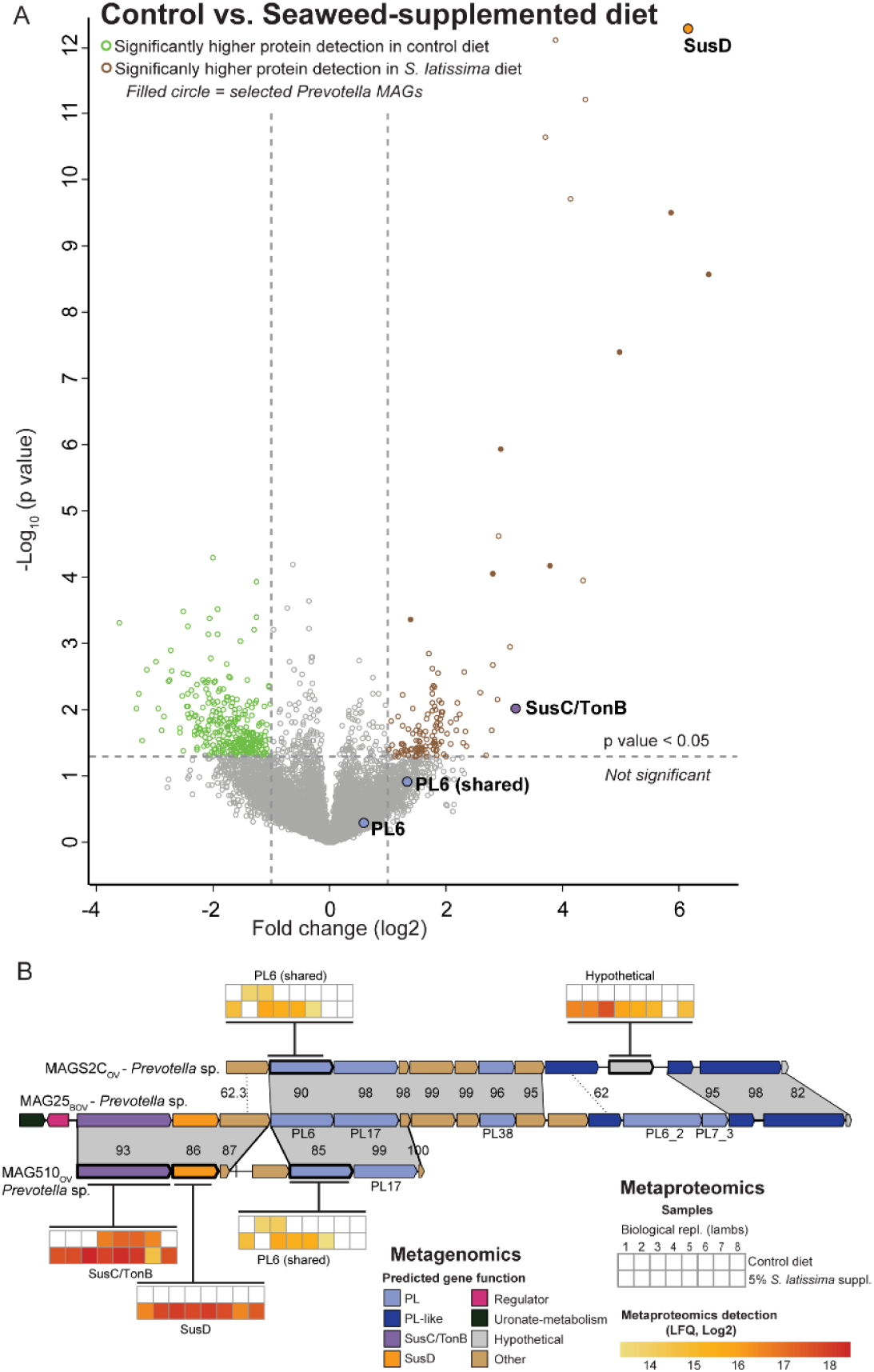
Detection of alginate lyases within AULs. A) Volcano plot of all protein groups identified in lamb rumen samples demonstrates detection of two alginate lyases (PL6) from *Prevotella* spp. (MAG510_OV_ and MAGS2C_OV_), along with a SusC and SusD that were significantly higher expressed in the 5% seaweed-supplemented diet group compared to the control group. B) According to the predicted gene organization, the PL6 enzymes and SusC/SusD were located within AULs that showed high synteny to AULs extracted from *Prevotella spp.* MAG25_BOV_ found in the cattle-based RUSITEC microbiome. Gray shading indicates homology and % identity values. Genes with bolded outlines were detected within the lamb metaproteome, and the heatmaps show the protein intensities (LFQ values for the eight replicates sampled in control and 5% *S. latissima*-fed lambs).

In addition to the detection of the PL6 enzyme, a proteome inspection of MAG510_OV_ also unveiled differential expression of a TonB-dependent receptor (SusC) and its adjacent SusD-like substrate-binding protein in the 5% *S. latissima* dietary group compared to the control. Specifically, the MAG510_OV_-affiliated SusD-like protein was the most upregulated protein with *S. latissima* supplementation in the entire dataset (**Figure 3A**). Another essential component of the Sus system, the SusE-like outer membrane protein, was also detected in the proteome of MAG510_OV_, yet at a lower LFQ detection level. Within a PUL, the *susC* and *susD*-like genes are typically situated in close proximity to the relevant CAZymes ^31^. However, due to genome fragmentation, the *susC*/*susD* pair and the alginate lyases (*pl6* and *pl17*) were found towards the edges on two separate contigs, and thus, a complete AUL could not be determined. Despite their physical separation on different contigs, possibly due to incomplete genome reconstruction, the explicit co-expression of *susC*/*susD* and *pl6* implies that these genes may still function as a part of a co-regulated AUL.

### *Prevotella* spp. AULs in globally separated rumen ecosystems

We observed that AULs found in *Bacteroidota* originating from the lamb rumen microbiome and cattle-based RUSITEC system from two different continents were highly syntenic and contained highly homologous sequences to the other AULs. In total, eight *Bacteroidota* phyla members had intact AULs, along with the AUC from the *B. bullata* MAG32_BOV_. All MAGs contained at least one PL6 and PL17 pair, with some containing either additional PL6, PL7, or PL38 sequences within the AULs (**Figure 4A**). Additionally, using Diamond Blast, AlphaFold ^32^, and Dali searches ^33^, uncharacterized AUL genes were identified as lyase-like sequences. Between RUSITEC and lamb *Bacteroidota* spp. MAGs, the highest homology was discovered between *Prevotella* MAGs, specifically the cattle-based RUSITEC MAG25_BOV_, and lamb rumen MAGs MAGS2C_OV_ and MAG510_OV_ showing amino acid homology as high as 99% (**Figure 3B, Extended Data 1**). This homology extends to the alginate lyases, as the PL17 member across the three AULs shared 98% amino acid identity, while the shared PL6 members exhibit 85% identity. The PL38 protein shared between MAG25_BOV_ and MAGS2C_OV_ displayed 96% sequence homology.

**Figure 4:**
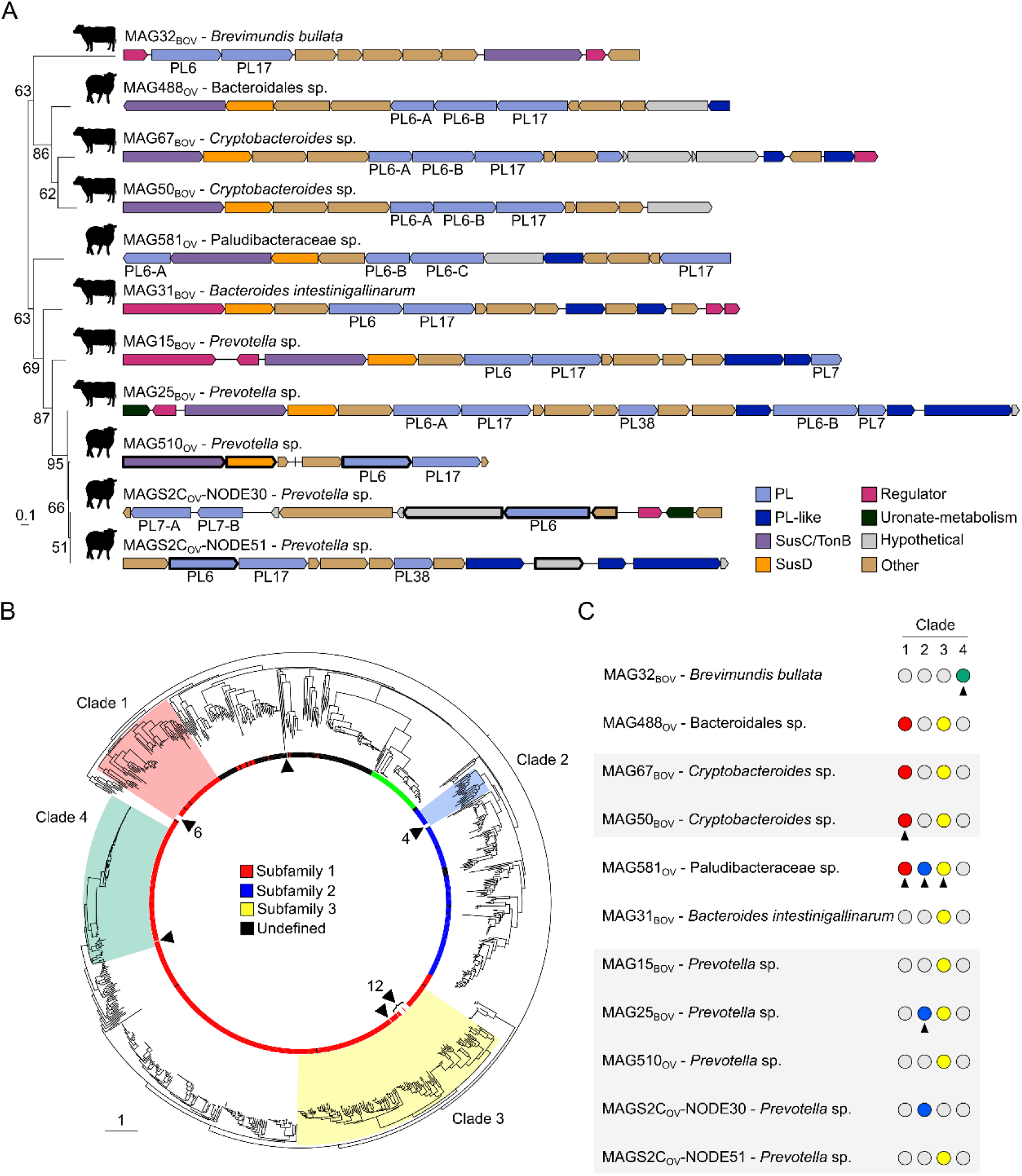
Synteny between lamb and cattle alginate AULs. A) Alginate AUL/AUCs from lamb and cattle. OrthoFinder was used to generate a species tree composed MAGs containing AUL/AUCs. The AULs and AUC were created with the gggenes package. Bolded genes were identified within the lamb metagenomics. B) Phylogenetic trees generated from all sequences within the PL6 CAZy database, including those predicted within lamb and cattle MAGs. The inner ring denotes CAZy subfamilies. Phylogenetic clades with uncharacterized MAG sequences are shaded and given labels 1-4, while MAG sequences are directly noted by black arrows and the number of sequences with the clade. C) Summary of which lamb and rumen MAGs containing AUL/AUCs contain PL6 members within clades 1-4. Black arrows indicate sequences which were selected for further characterization.

Using SACCHARIS 2.0 ^85^, the PL6 enzymes within *Bacteroidota* AULs, along with homologous PL6s found on shorter contigs in both the RUSITEC and lamb MAGs, partitioned into three distinct clades based on amino acid homology. In addition, the PL6 enzyme found within *B. bullata* MAG32_BOV_ formed a more distantly related group (Clade 4) compared to the other three clades. Aligning these PL6 members to all PL6 enzymes within the CAZy database revealed that Clade 1, 3, and 4 consisted entirely of PL6 subfamily 1 members, while Clade 2 was composed of subfamily 2 members (**Figure 4B**). Further inspection of the phylogeny showed that PL6 members of both Clade 1 and Clade 2 were most closely related to characterized sequences from *Rhodothermus marinus* DSM 4252. Specifically, Clade 1 members shared the highest sequence similarity (37-40% identity) with Rmar_1165, an enzyme active on poly-MG. Similarly, Clade 2 members were most closely related to Rmar_1386 (38-42%), also active on poly MG (**Supplementary Figure S4**) ^34^. A majority of Clade 3 members had the highest sequence similarity (41-48%) to an exo-type alginate lyase of human gut bacterium *Bacteroides clarus* ^35^, except MAG581_OV_-C, which was more similar (33.43%) to a poly-MG-active PL6 enzyme characterized from the marine bacterium *Nonlabens ulvanivorans* ^34^. Lastly, the MAG32_BOV_ PL6 member shared 55% sequence similarity with an endo-poly-MG active PL6 from *Stenotrophomas maltophilia* KJ-2 (**Supplementary Figure S5**) ^36^.

Overall, PL6 members of Clade 3 were found in all *Bacteroidota* MAGs encoding AULs, but not within the AUC identified in MAG32_BOV_. Near half of the PULs contained multiple PL6 members belonging to different clades, of which the combination of Clade 1 and Clade 3 were most prevalently occurring in both lamb rumen (*Bacteroidales* sp. MAG488_OV_ and *Paludibacteraceae* sp. MAG581_OV_) and cattle-based RUSITEC MAGs (*Crypobacteriales* spp. MAG50_BOV_ and MAG67_BOV_) (**Figure 4C**). The *Prevotella* MAGs seemed to rather combine PL6 Clade 3 with Clade 2 within the same AUL (MAG25_BOV_ and MAG581_OV_), or at separate loci (MAGS2C_OV_). Of note, the two *Prevotella*-affiliated PL6 proteins detected in the metaproteome belonged to Clade 2 and Clade 3 (shared PL6).

To confirm the predicted alginate activity of AUL/AUCs, PL6 members from each clade (Clade 1: MAG581_OV_-A, MAG50_BOV_-A; Clade 2: MAG581_OV_-B, MAG25_BOV_-B; Clade 3: MAG581_OV_-C; Clade 4: MAG32_BOV_) were selected for gene synthesis and functional characterization. All selected PL6 members contained SPI signal peptides, except for MAG50-A which had an SPII signal peptide (not shown, as signal peptides were ultimately removed for protein expression in this study). Enzymes were incubated with alginate from brown seaweed and observed after overnight digestion, as well as continuous direct measurement of unsaturated product formation at 232 nm. Direct measurement was used to determine the optimum pH and initial velocities of the selected PL6 enzymes in the linear range of the reaction (**Supplementary Figure S6**). All PL6 members, except MAG581-B, produced absorbance at 232 nm, indicating enzymatic activity. However, kinetics of most PL6 enzymes demonstrated inefficient digestion of alginate, with the exception of MAG50_BOV_-A, and Michaelis-Menten could not be accurately fitted (**Supplementary Figure S6F**). Despite variations in catalytic efficiencies, all PL6 members produced oligosaccharides when incubated with alginate (**Figure 5A**). Using HPAEC-PAD, PL6 members from the same clade displayed overlapping activities. LC-MS was used to further characterize the structure of unsaturated oligosaccharides (**Figure 5B and C**).

**Figure 5:**
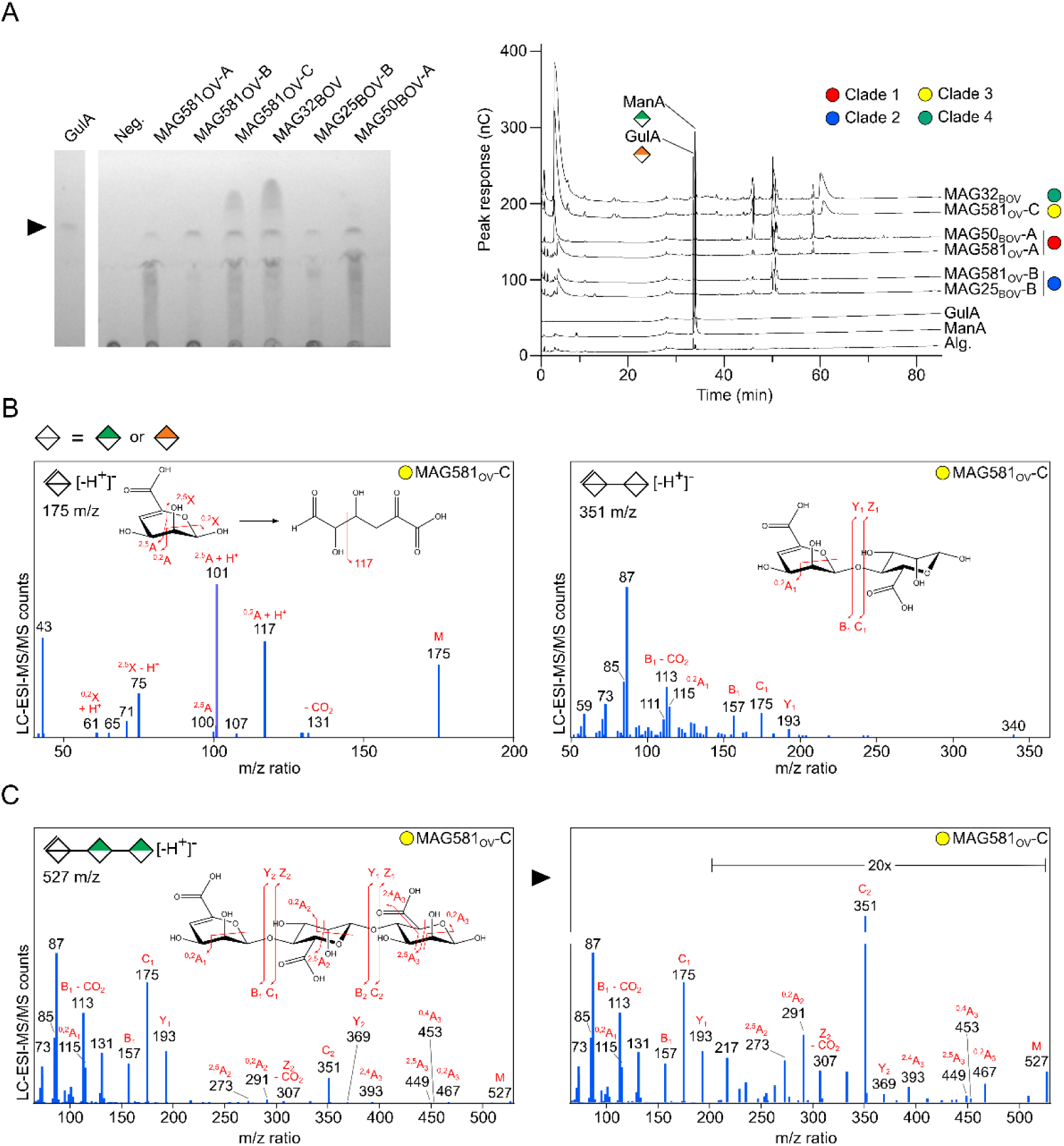
Characterization of alginate oligosaccharide products from *Bacteroidota* species PL6 alginate lyases. A) TLC (left) and HPAEC-PAD (right) product analysis of brown seaweed alginate digested with lamb and cattle-associated MAG PL6 enzymes. B) Alginate lyase oligosaccharide products were analyzed by LC-ESI-MS/MS. Ions of m/z 175 (left) and 351 (right) corresponding to unsaturated mono- and disaccharide products were observed, and spectra were extracted. ESI-MS/MS with HCD was performed in order to confirm identity of the extracted ions, and MS2 product ion spectra are shown with the carbohydrate fragmentation depicted and identified. C) MS2 spectra of ion 527, corresponding to unsaturated trisaccharide, identified as ΔHexA-ManA-ManA based on the presence of signature ions ^40^. Monosaccharide symbols are displayed according to the Symbol Nomenclature for Glycans system ^41^. For monosaccharides unable to be discerned between mannuronate and guluronate by LC-MS, white symbols are used.

All PL6s produced alginate oligosaccharides of varying degrees of depolymerization. Clade 3 and Clade 4 members produced monosaccharides that appear to decompose through a keto-enol intermediate into 4-deoxy-L-erythro-5-hexoseulose uronate (DEH) ^37, 38, 39^. Both MAG581_OV_-B and MAG25_BOV_-B appeared less active on brown seaweed alginate, producing only small amounts of alginate oligosaccharides.

### Direct Interactions between *S. latissima* polysaccharides and microbial cells

The capability of the RUSITEC microbial communities to interact with *S. latissima* polysaccharides was investigated *in vivo* using fluorescent *S. latissima* extract (FLA-SLAT). Cells within the 2% and 50% *S. latissima* treated RUSITEC systems demonstrated uptake of FLA-SLAT, which overlayed with DAPI staining (**Figure 6A**). Total cell density at the beginning of the RUSITEC and after 15 days was enumerated by the number of DAPI-stained cells. The RUSITECs started with an average of 4.2×10^-7^ cells ml^-1^ and showed the same abundance at day 15. Furthermore, 1 h and 1 day incubations with the FLA-SLAT probe were performed with the control and 15 day RUSITECs and these values were compared to total cell count. Within the control RUSITEC communities, under 7.0% of the RUSITEC starting fluid community took up FLA-SLAT after 1 h and 1 day, whereas the control diet jumped from 3.9% at 1 h to 13.3% after 1 day (**Figure 6B**). This is in contrast to the 2% and 50% *S. latissima* communities, where 26.8% and 16.2% of the community took up FLA-SLAT after 1 h of incubation, which then decreased to 7.1% and 14.8%. SR-SIM images further showed that cells which retained the FLA-SLAT probe concentrated the probe nearest to the outer membrane of cells (**Figure 6C**). Uptake of fluorescent alginate (FLA-ALG) by *S. latissima* treated RUSITEC samples was also observed (**Supplementary Figure S8**). The 2% *S. latissima* treatment showed a more rapid uptake of FLA-ALG at 1 h compared to the 50% *S. latissima* treatment at 1 day.

**Figure 6:**
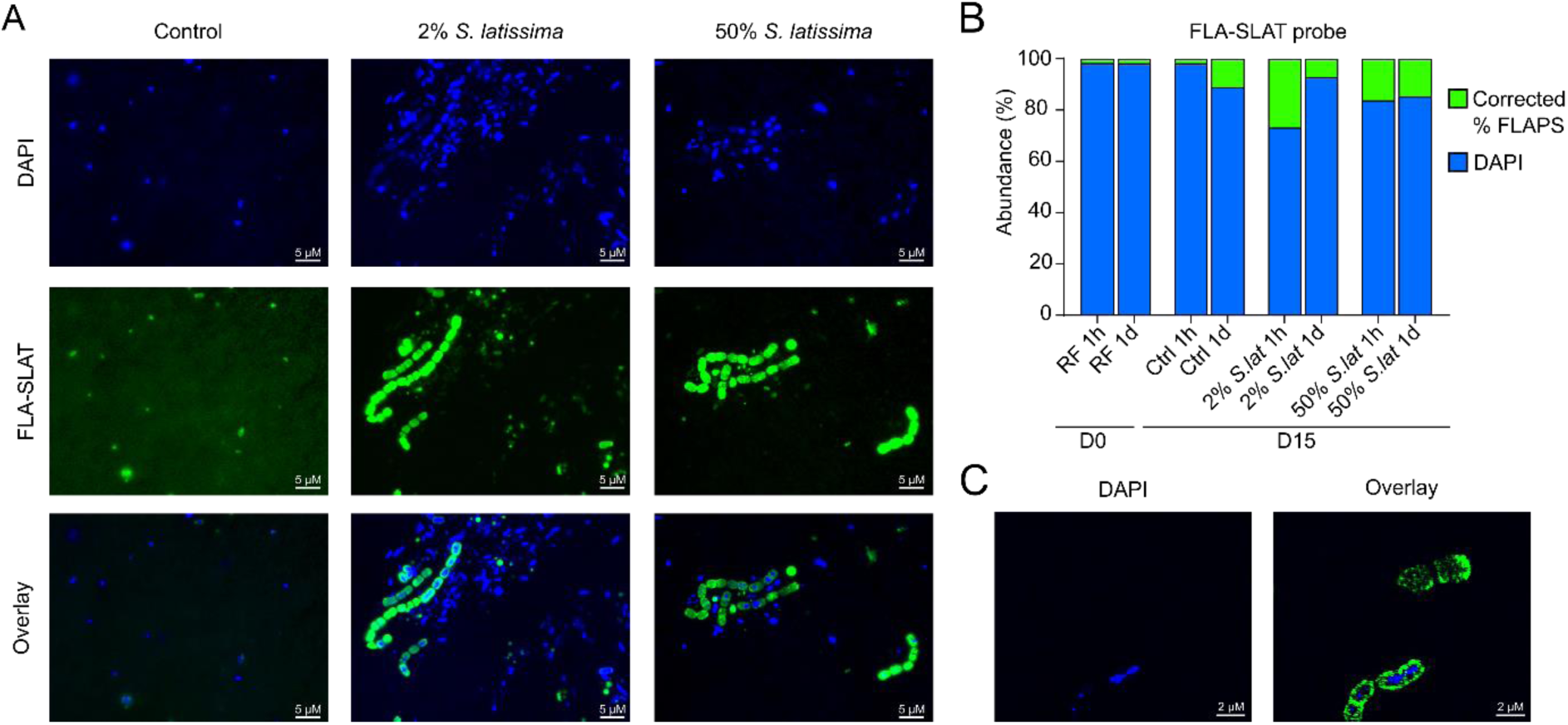
Epifluorescence visualization of FLA-SLAT interactions in RUSITEC rumen microbial communities that were enriched with 50% *S. latissima* for 15 days. A) FLA-SLAT interactions in control, 2% *S. latissima*, and 50% *S. latissima* RUSITEC microbial communities sampled from RUSITEC vessels after 15 days, stained with DAPI (blue) and incubated with 0.2 % FLA-SLAT (green) for 1 day. B) Relative abundance of the enumerated microbial communities for D0 and D15 RUSITEC sample after incubation with FLA-SLAT after 1 h and 1 day, where cells that interacted with the FLA-SLAT probe (green) and cells that only displayed a DAPI signal (blue). RF – RUSITEC fluid, Ctrl – control, Corrected % FLAPS – autofluorescence subtracted. C) SR-SIM of cells from 50% *S. latissima* experiment after 1 day FLA-SLAT incubation. Cells were stained by DAPI, FLA-SLAT, and an overlay of DAPI and FLA-SLAT is shown.

## DISCUSSION

### *S. latissima* cell wall structure

The matrix cell wall polysaccharides of brown seaweeds are largely comprised of anionic polysaccharides, such as alginate and fucoidan ^6^. Their charge density corresponds with the ionic nature of ocean water and enables metal-dependent, higher-order polymer cell wall dynamics, such as gelation. The polysaccharide composition of *S. latissima* was confirmed to consist of cellulose and mixed-linkage glucans, and the matrix polysaccharides fucoidan and alginate. Alginate comprised approximately 15% of the total plant cell wall polysaccharides in S*. latissima* (**Figure 1D; Supplementary Table S1**), which is in agreement with previous reports ^24^. Fucoidan comprised ∼34% of the estimated polysaccharide content, more than two-fold that of alginate. *S. latissima* fucoidan is known to be structurally complex, consisting of highly sulfated α-1,3-L-fucan backbones, with side chains composed of β-1,6-D-galactose or alternating β-D-glucuronic acid and α-D-mannuronic acid extensions (Bilan et al., 2010). Despite the abundance of fucoidan, we were unable to detect any putative fucanases from the lamb feeding experiment or bovine RUSITEC metagenomic and metaproteomics datasets, suggesting that fucoidan is likely not digested by either ruminant. To date, the discovery of a fucanase or fucoidan-specific sulfatase from a terrestrial gut microorganism has not been reported. Whether this stems from its structural complexity, bioavailability, or other factors remains to be determined.

### Structure and abundance of AULs from ruminant datasets

*Bacteroidota* MAGs with the genetic capacity to degrade alginate were constructed from the lamb feeding experiment and bovine RUSITEC sequencing data. Intriguingly, MAGs that encoded the capacity to consume alginate included two closely related *Prevotella* strains that exhibited diet-induced dose responses in their relative abundance (**Figure 4**). Metaproteomics further confirmed these *Prevotella* species produced polysaccharide lyases specific for the digestion of alginate (**Figure 3**). Reconstruction of the pathway architecture containing these alginate lyases indicated they were functional components of AULs, similar to those previously observed in marine and human gut bacteria ^13, 16, 42^ (**Figure 4**). Furthermore, the detection of elevated levels of SusC-like and SusD-like proteins in the *S. latissima* diet group reinforces the relevance of these AULs for alginate digestion. Surprisingly, a large-scale diet-induced difference was not observed in the overall microbial community structure when evaluated by 16S rRNA gene sequencing. This suggested that a limited amount of specific rumen microbial species play central roles by occupying a specialized niche for digestion of seaweed-derived polysaccharides. Notably, other PL enzymes associated with alginate consumption were also identified within the AULs, though their protein expression was not detected. This could be due to several factors, such as inclusion level, compartmentalized secretion, specificity of induction products ^43, 44^, or insufficient adaptation time for efficient utilization of seaweed. Others have suggested that alginate polymers decompose slowly ^42, 45^.

Increasing the overall alginate concentration to 50% of the total mixed ration (25-fold over the lamb feeding experiment) within the RUSITEC experiment, allowed us to discern multiple *Bacteroidota* organisms capable of digesting alginate (**Figure 4A**). In contrast, *Prevotella* spp. abundance was not significantly different between control and *S. latissima*-supplemented RUSITECs, and, although expected ^46^, we identified multiple *Prevotella* spp. within RUSITEC-derived MAGs containing AULs.

To measure direct interactions between polysaccharides and microorganisms in the rumen, we previously developed a technique using fluorescent polysaccharides, or FLAPS ^47^. This enables the response of microorganisms to labeled substrates to be quantified at the single-cell level. Accumulation of fluorescent polysaccharide extracts from *S. latissima* (FLA-SLAT) (**Figure 6**) and alginate (FLA-ALG) (**Supplementary Figure S8**) in microbial cells, provided further support that alginate is consumed and this occurs in response to incubation with *S. latissima*. SR-SIM imaging following FLA-SLAT incubation showed that most interactions with FLA-SLAT occurred in the periphery of cells, which is reminiscent of Gram-negative bacteria that employ a selfish mechanism of foraging ^47, 48, 49^.

### Function of PLs from ruminant AULs

The operonic AUL structures within this study were identified using previously characterized lyase families: PL6, PL7, PL17, and PL38. While members of PL6 have been shown to be both endo- and exo-acting, they typically display an endo-mode of action, indiscriminately cleaving internal glycosidic bonds within the alginate polymer to produce oligosaccharides with unsaturations at their non-reducing ends. In contrast, PL17 enzymes are often categorized as exo-acting oligoalginate lyases that primarily act on the polyM structure at the end of an alginate chain, releasing unsaturated mono- or disaccharides of mannuronate ^13^. Previous studies of alginate consuming marine and human gut bacteria have shown that PL6 and PL17 members are frequently co-located, strongly suggesting that they may operate in a coordinated manner to efficiently depolymerize alginate ^13, 16^. In addition to PL6 and PL17, some MAGs encoded alginate lyases belonging to the PL7 and PL38 families. To confirm the function of representative enzymes from these AULs, several PL6 members were selected for biochemical characterization. PL6 enzymes can possess “vanguard” status by catalyzing the first stages of alginate depolymerization at the cell surface ^50^. Several recombinant PL6 enzymes were shown to produce substantial amounts of unsaturated products by TLC, HPAEC-PAD, and LC-MS. Due to the complexity of M/G orientation and ratio, and the ability of PL6 enzymes to change mode of action (endo/exo) depending on either alginate substrate (poly-ManA or poly-GulA) ^34^, and the absence or presence of non-catalytic domains ^35^, precise activity was not demonstrated within this study. Notably, delineation between ManA and GulA signatures are often indistinguishable using methods other than NMR; however, prediction of some oligosaccharide structures is possible based upon a previous study that defined patterns of differential MS2 fragmentation abundance of alginate oligosaccharides. For example, an identified unsaturated tri-saccharide, was found to contain signature ions (m/z of 307, and high ratio of 449/453) suggesting the oligosaccharide product consists of a ΔHexA-ManA-ManA ^40^ (**Figure 5C**). Further, EI-MS and MS2 fragmentation revealed that some uronic acid residues of alginate may in fact be sulfated (**Supplementary Table S7**). This unexpected result warrants further study.

### Evolution of alginate metabolism within the rumen microbiome

Seaweed polysaccharides tend to be structurally and chemically complex, highly charged, and absent from terrestrial plants typically consumed by livestock. Members of the human microbiome are known to have catabolic pathways dedicated to digestion of polysaccharides from red ^14, 17, 18^ and brown ^13^ seaweeds. Multiple hypotheses, including the ‘sushi factor’ ^15, 17^ and ancient transfer ^13^, have been proposed to explain the origin of these pathways in gut microbiomes. In contrast to humans, much less is known about the mechanisms by which livestock microbiomes adapt to the introduction of exotic dietary polysaccharides, such as alginate from brown seaweed. Previously, alginate-consuming *Prevotella* spp. were isolated from the North Ronaldsay sheep that live on the Orkney Islands, which have a diet augmented with brown seaweeds, likely *Laminaria*, *Fucus,* and *Ascophyllum* species ^22^. How these animals survived a radical change in their diet and lifestyle, and the evolutionary processes that drove the microbial adaptation towards seaweed polysaccharide catabolism has not been thoroughly studied. Recent studies have also revealed that wild Svalbard reindeer populations are adapting to the warming Arctic climate by supplementing their diet with kelp, driven by an increasing prevalence of ice-locked pastures ^51^.

Here, we discover and characterize the structure of syntenic AULs responsible for alginate catabolism from both cattle RUSITEC and lamb microbial communities exposed to dietary *S. latissima*. Alginate catabolism appears to be a core functional trait performed by autochthonous members of the ruminant microbiome ^52^, indicating that these genomic loci were not recently transferred from the surface microbiome of *S. latissima* or another marine ecosystem. The conservation of PL6 and PL17 pairs is shared within *Bacteroidota* spp. from geographically and taxonomically distinct ruminants, suggests that these genetic loci were acquired by one or a series of ancient horizontal gene transfer events, similar to what was described from humans ^13^. In this light, perhaps AULs may have evolved in an ancestor common to humans and cattle. Despite conservation in function, there is noticeable structural variation in the gene content and organization (**Figure 4**). PL6 enzymes from Clades 1 and 2 were most closely related to PL6s from a soil and marine bacterium respectively, and MAG581-C in Clade 3 was most closely related to a PL6 member in marine *Dysgonomonas alginatilytica*. This suggests that these enzymes likely originated by transfer from a marine bacterium (**Supplementary Figure S4**). Regardless of how individual pathways assembled, the overall functional conservation of AULs within the ruminant microbiomes is clearly maintained throughout lineages and between populations. In this manner, the maintenance of alginate catabolism in the absence of substrate availability represents a “latent trait” within the rumen microbiome. Often, these pathways are encoded within the genomes of saccharolytic generalists, which likely provides a mechanism for retention of alginate competent strains when these substrates become scarce. Ultimately, this provides a nutritional benefit to the host. In this regard, acquisition and maintenance of a diverse metabolic capacity appears to position the animal to rapidly adapt to abrupt changes in its diet and provides tantalizing clues into the ancient foraging habits of its ancestors.

## CONCLUSION

The adaptation of livestock to novel feedstocks may provide a solution to pressures associated with climate change and land use competition with food crops. Seaweed mariculture represents a source of novel feed ingredients for ruminant livestock; however, how and at what rate ruminant microbiomes can adapt to the introduction of new carbon sources may dictate the overall feasibility of their incorporation into production systems. Here we investigated the ability of two geographically and taxonomically distinct ruminant microbiomes to adapt to *S. latissima*, a widely distributed, commercially-grown brown seaweed. Our study demonstrates that rumen *Bacteroidota* spp. contain highly conserved AULs that are upregulated in *S. latissima* supplemented diets. Further, these PULs appear to be native to the ruminant microbiome, not the marine environment, and capable of digesting dietary alginate. This opens up new hypotheses about the ability of ruminant microorganisms to adapt to other exotic polysaccharides and novel forages by the retention of latent traits that are carried within the microbiome of ruminant livestock.

## MATERIALS AND METHODS

### Preparation of unfractionated cell wall of *S. latissima*

*S. latissima* was harvested from the waters near Vancouver Island, B.C., Canada on May 1^st^, 2022. The cell wall extractions were prepared in accordance to Avci, Pattathil ^53^ with modifications according to previous reports ^24, 54^. Approximately 100 mg of ball-milled dry sample was extracted in 40 mL of absolute ethanol (EtOH) for 8 h, followed by three 8 h extractions in 40 mL of 80% (v/v) EtOH, and then two 20 min washes with absolute EtOH. After each alcohol extraction and wash, samples were centrifuged at 3,000 × *g* for 30 min. The final alcohol insoluble residue (AIR) was vacuum-dried by SpeedVac (Thermo Scientific, USA).

### Glycosidic linkage analysis of cell wall

The AIR of *S. latissima* was analyzed as described ^24^. Briefly, polysaccharides in dry AIR power (10 mg) were methylated by 1.2 mL of methyl iodide in 2 mL of dimethyl sulfoxide in the presence of excess sodium hydroxide ^55, 56^. The methylation product was suspended in dichloromethane (DCM) and partitioned against 10% acetic acid in water (v/v) over ice once then against deionized water three times. After each partitioning, the upper phase was collected and pooled. Following the final partitioning, the lower phase was evaporated to dryness, and the pooled upper phase was mixed with the evaporated lower phase. The mixture was dialyzed with a molecular weight cut-off of 6,000 to 8,000 Da against 0.1 M triethylamine hydrochloride (TEAH) deionized water solution then against pure deionized water, followed by freeze-drying. Two additional rounds of methylation were conducted with the 0.1 M TEAH dialysis step omitted ^24^. The permethylated sample was converted to partially methylated alditol acetates (PMAAs) by hydrolysis in 2 mL of 4 M trifluoroacetic acid (TFA) at 100 °C for 4 h, reduction with 20 mg of sodium borodeuteride (NaBD_4_, 99 atom % D, Alfa Aesar) in 2 mL of deionized water for 16 h ^57^, and acetylation by heating in a mixture of acetic anhydride and TFA (5:1, v/v) at 60 °C for 1 h ^24^. In a separate experiment, the dry AIR powder (10 mg) was subjected to weak methanolysis (0.5 M methanolic HCl, 80 °C, 20 min), followed by NaBD_4_ reduction to convert uronic acids to their corresponding 6,6’-dideuterated neutral sugars ^58, 59, 60^. Excess NaBD_4_ was neutralized by acetic acid, followed by evaporation of the product to dryness and repeated evaporation in 10% acetic acid in methanol (v/v) then in absolute methanol to remove borate, before the sample was converted to PMAAs ^24^. All PMAAs were analyzed using an Agilent 7890A-5977B GC-MS system (Agilent Technologies, Santa Clara, CA, USA) coupled to a medium-polarity Supelco SP-2380 column (60 m × 0.25 mm × 0.2 µm; Sigma-Aldrich, USA) under a consistent flow of helium at 0.8 mL/min. Inlet temperature was maintained at 250 °C. Sample was injected with a 10:1 split ratio. The oven program started at 120 °C (hold 1 min) and then increased at 3 °C/min to 200 °C (hold 50 min) then at 3 °C min/min to 250 °C (hold 20 min). The PMAAs were identified by comparing their EI-MS fragmentation patterns and retention times to those of prepared PMAA standards ^24^ and also by referencing the literature (Carpita and Shea, 1989). Relative molar linkage compositions of the PMAAs were determined from the TIC chromatogram, following the principle that a PMAA’s quantity correlates with the ratio of its TIC peak area to its molecular weight, as described in the published protocol ^61^. The peak corresponding to the PMAA of 4-Glcp was used to normalize the relative abundances of neutral sugar PMAAs in the untreated sample and the PMAAs derived from uronic acid linkages in the sample pretreated with weak methanolysis-NaBD₄ reduction ^24^. Three separate experiments were conducted to the AIR sample.

### *In vivo* experimental design

A total of 24 Norwegian White lambs were used in the *in vivo* feeding experiment, as previously explained in detail by Grabež, Devle ^62^. In brief, weaned ewe lambs with body weight 37.3 ± 1.6 kg were randomly assigned to three dietary treatment groups (n = 8 per group). All lambs had free access to clean drinking water and were fed the experimental diet ad-libitum twice a day (at 08:00 h and 14:00 h) in individual pens. The control group (no seaweed included) were fed a total mixed ration of grass silage and compound feed, rolled barley and mineral premix. The two other treatment groups were additionally supplemented with 2.5% and 5.0% *S. latissima* on a dry matter (DM) basis. The *S. latissima* used in the *in vivo* feeding experiment was harvested by Seaweed Solution AS (Trondheim, Norway) at their cultivation site outside Frøya, Norway (collected 28/05/2018), and immediately frozen. Prior the feeding experiment, the seaweed biomass was chopped to uniform particle size and wilted, before stored frozen again until final preparation of experimental diets. The experimental period lasted for 35 days, during which the lamb growth rate, daily feed intake and feed dietary composition were regularly monitored. All animal procedures were approved by the committee overseeing the rules and regulations governing animal experiments in Norway under the surveillance of the Norwegian Food Safety Authority (FOTS-ID: 16406).

### RUSITEC experimental design

The RUSITEC experiment followed a 2 x 2 + 1 factorial design with two experimental runs and two treatments. Each RUSITEC unit was equipped with eight 920 mL fermenters, designated with three replicates per treatment (n = 6) in each unit across both runs. Here, the basal substrate consisted of a 50:50 (DM basis) barley silage and barley straw diet. The two dietary treatments included control (no seaweed included) and *S. latissima* (collected from Vancouver Island, B.C., Canada, 01/05/2022) included at 2% dietary DM. An additional fermenter in each run was fed 50% *S. latissima* for microbial profiling. The seaweed replaced equal proportions of barley silage and barley straw, and treated was randomly allocated to a RUSITEC fermenter in each unit. Prior to the experiment, the barley straw, barley silage, and seaweeds were dried at 55 °C for 48 h and then ground through a 4 mm screen using a Wiley Mill (standard model 4; Arthur H. Thomas Co., Philadelphia, PA, USA). A total of 10 g DM was included in nylon bags (R1020; 10 × 20 cm; 50 ± 10 μM porosity; 107 R1020, ANKOM Technology, Macedon, NY, USA) for incubation within the two RUSITEC units.

Rumen inoculum was obtained from three ruminally cannulated beef heifers previously adapted to a barley silage and barley straw-based diet. Donor heifers used in this experiment were cared for in accordance with the guidelines of the Canadian Council on Animal Care (2009) and were approved by the Institutional Animal Care and Use Committee (ACC2304). The rumen contents were collected 2 h pre-feeding from four different sites within the rumen and squeezed through PECAP mesh (mesh size 250 µm; PA66CG-250 136 cm, Sefar Nytal, Gilbert Saguenay, QC, CA). Both the solid and liquid proportions were pooled in equal proportions from each heifer, and transported to the laboratory in an insulated thermos kept at 39 °C. All fermenters were maintained at 39 °C in a water bath before filling with 180 mL of pre-warmed McDougall’s buffer (McDougall, 1948), and 720 mL of double strained rumen fluid added under a stream of CO_2_ to maintain anaerobic conditions. One R1020 Ankom bag filled with 20 g (wet weight) of mixed rumen solids, and one bag with the dietary treatment were added to each fermenter. The fermenters were then fitted within each RUSITEC unit contained within a circulating water bath retained at 39 °C. After 24 h, the bag containing the rumen solids was replaced with a dietary treatment bag, such that each day the bag incubated for 48 h was replaced with a new bag. Artificial saliva was continuously infused into fermenters (26 mL h^-1^) using a peristaltic pump set to achieve a dilution rate of 2.9% h^-1^. The effluent and gas were collected within 2 L Erlenmeyer flasks, and 4 L reusable gas tight collection bags Curity®; Conviden Ltd., Mansfield, MA, USA), respectively, connected through a closed tubing system. The RUSITEC experimental period lasted for 15 days, with day 1-7 used for adaptation and day 8-15 used for sampling and measurements. The DM, OM, CP, and NDF degradability, and total VFA concentration was determined from day 8-15 and analysis was conducted as previously described (Terry et al., 2023).

### Microbial sampling and DNA extraction

Samples for microbial analysis were collected from both the RUSITEC experiment and the *in vivo* feeding trial. These samples were subsequently used for 16S rRNA and shotgun metagenomics.

From each RUSITEC fermenter, 2 mL of liquor was collected at day 0, 1, 8 and 15. Samples were centrifuged at 20,000 x *g* for 20 min and supernatants were removed. Pellets were suspended in 1 mL of DNA/RNA Shield™ (Zymo Research, USA) before genomic extractions were performed using the DNeasy PowerSoil Pro kit (Qiagen, Canada) following the manufacturers protocols. DNA was stored at −80 °C until analysis.

In the *in vivo* feeding trial, rumen samples were collected at the end point of the experiment period. Here, all animals were slaughtered at a commercial slaughterhouse (Rudshøgda, Nortura SA, Norway) and their intact gastrointestinal tracts were directly moved to a working bench. The stomach was opened, and the reticulo-rumen content was hand-mixed prior to sampling. The mixed sample was transferred into a sterile strainer blender bag (0.50 mm pore size; Stomacher® 400 Seward BA 6041, Worthing, UK) and gently squeezed to separate the fluid and particle phases. The phases were then sampled into cryotubes, transported in liquid nitrogen to the laboratory and stored at − 80 °C until analysis. Upon microbiome analysis, samples from both fluid and particle phase were thawed on ice and homogenized by vortexing prior cell lysis and DNA extraction. For cell lysis, 0.25 g homogenized biomass was weighted out for bead-beating using a FastPrep-24 Homogenizer (MP Biomedicals LLC., Ohio, USA) at 4m/s for 45 seconds. DNA were extracted using DNeasy PowerLyzer PowerSoil kit (Qiagen, Hilden, Germany) following to the manufacturers protocol. The extracted DNA were quantified using a Qubit Fluorimeter and the Qubit dsDNA BR Assay Kit (ThermoFisher Scientific, Waltham, MA, USA) and stored at −80 °C until further use.

### 16S rRNA gene sequencing

In the RUSITEC experiment, the purified DNA samples were sent to Génome Québec (Montréal, QC, Canada) for Illumina MiSeq PE250 16S rRNA gene sequencing using the primers: 515F (5’-GTGCCAGCMGCCGCGGTAA-3’) and 806R (5’-GGACTACHVGGGTWTCTAAT-3’) targeting the V4 region of the 16S rRNA gene. Paired end reads were quality trimmed using Trimmomatic ^63^ with a sliding window of 5:20, and adapters were automatically removed. Trimmed fasta files were merged and classified using Kraken 2 ^64^ according to the Silva 138.1 SSU database. Bracken ^65^ was used to estimate bacterial and archaeal abundance at the genus level.

Liquid and particle phase samples from all 24 lambs was collected for the amplification of the 16S rRNA gene. The 16S rRNA gene was PCR-amplified using Pro341F/Pro805R primer pair ^66^, which targets the V3-V4 region of both bacteria and archaea. The 25 μL PCR reactions consisted of 1× iProof High-Fidelity Master Mix (Bio-Rad, Hercules, CA, USA), 2.5 μL each primer, and 7.5 μL template DNA. PCR thermal cycling began with a hot start step at 98 °C for 3 minutes, and they were followed by 25 cycles of 98 °C denaturation for 30 s, 53 °C annealing for 30 s and 72 °C extension for 30 s, followed by a final extension step at 72 °C. The amplicon libraries were purified with AMPure XP beads (Beckman Coulter, Indianapolis, IN, USA) and indexed with the Nextera XT Index Kit v2 (Illumina, San Diego, CA, USA). Equimolar concentrations of the libraries were pooled together, and further purified with AMPure XP beads. The pooled library was quantified using Qubit Fluorometer, diluted and denatured before sequenced on an Illumina MiSeq sequencing system using 2 x 300 bp cycle runs with the MiSeq reagent kit v3. The generated 16S rRNA gene sequences were processed using the DADA2 pipeline ^67^ in R v4.3.2 ^68^ which included quality control, elimination of primers and adapters. The sequences were clustered into Amplicon Sequence Variance (ASVs) and the ASV identified were taxonomically assigned using SILVA Database release 138.1 ^69^. Downstream analysis of lamb and RUSITEC communities are described in detail in **Supplementary Text S2**.

### Metagenomics

To reconstruct a core rumen microbial genome catalogue from each experimental group of lambs, extracted DNA from corresponding fluid and particle from four animals in each diet group (in total 24 samples) were subjected for metagenome sequencing. The selection of these representative samples was guided by diversity and abundance metrics from the 16S rRNA gene analysis. Shotgun metagenomic sequencing was performed at the Norwegian sequencing center (Oslo, Norway), where the libraries were prepared using PCR-free TruSeq chemistry and sequenced on two lanes of the Illumina HiSeq6000 to generate 2 x 150 bp pair-ended reads. From the RUSITEC system, DNA from samples collected on experimental day 15 was sent for shotgun metagenomics at Génome Québec (Montréal, QC, Canada). These samples were sequenced on an Illumina NovaSeq6000 platform, also generating 2 x 150 bp pair-ended reads.

### Generation of metagenome-assembled genomes (MAGs)

RUSITEC sequences were trimmed with trimmomatic v0.39 and assembled using metaSPAdes v3.13.0. Assembled contigs were binned using the MetaWRAP v1.3.2 ^70^ binning and bin-refinement pipeline, using CONCOCT v1.1.0 ^71^, Maxbin2 v2.2.6, and Metabat2 v2.12.1. Bins were dereplicated using dRep v3.2.2 and filtered for 50% completion and under 10% contamination. CheckM v1.1.3 was used to evaluate the quality, while MAG abundance was estimated using CoverM v0.6.1. MAGs were annotated using dbCAN v3.0.7 ^72^ and Bakta v1.9.3^73^. GTDB-Tk v2.4.0 was used to infer taxonomic classification.

All metagenome raw reads from lamb samples were quality filtered using trimmomatic v0.36 ^63^ in pair end mode before the trimmed reads were assembled into contigs. Both individual assemblies and co-assemblies of all samples were carried out metaSPAdes v3.13.0 ^74^ and MegaHIT v1.2.9 ^75^, respectively. Contigs from the single assembly were binned using VAMB v3.0.2 ^76^, while the contigs from the co-assembly was binned using both MetaBAT2 v2.12.1 ^77^ and MaxBin2 v2.2.7 ^78^. Collectively, these three binning strategies generated 1,532 bins across all samples. These bins were re-dereplicated at 99% ANI using dRep v3.2.2 ^79^, which resulted in 291 dereplicated bins, hereafter referred to as metagenome-assembled genomes (MAGs), with completeness above 50% and contamination below 25%. This collection of MAGs were used as genome database for metagenome-centric metaproteome analysis, allowing to retain genetic information without compromising the sensitivity and false discovery rate estimation^80^. For biological interpretation, only MAGs with contamination > 10% were considered. CheckM v1.1.3 ^81^ was used to evaluate the quality parameters of each MAG, while CoverM v0.6.1 (https://github.com/wwood/CoverM) in genome mode was used to estimate the MAG abundance in each sample. Taxonomically classification of the MAGs was assigned using GTDB-Tk v2.4.0 ^82^, and function annotation carried out using dbCAN, PFAM, and KEGG, integrated in the DRAM v1.2.4 ^83^ annotation tool.

AULs from lamb and RUSITEC data were manually screened for SusC and SusD pairings. Syntenic AULs were compared using Diamond BLASTp v2.1.8 ^84^ and alignments were visualized using RStudio and the gggenes package ^85^. MAG homology was completed using OrthoFinder v2.5.5 ^86^, and CAZyme phylogeny was completed using SACCHARIS 2.0 v2.0.0.dev19 ^87^.

### Metaproteomics

#### Protein extraction and sequencing

Sample preparation for metaproteomic analysis was carried out using the particle phase of rumen samples collected from all lambs in the control and 5% *S. latissima* experimental group. Lysis buffer (10 mM DTT, 100 mM Tris-HCl (pH=8) and 4% SDS) and 4 mm glass beads (≤ 160 μm) were added to 500 μL of rumen samples, followed by brief mix and resting on ice for 30 minutes. Mechanical cell lyses was performed using FastPrep-24 Classic Grinder MP Biomedical, Ohio, USA) for 3 × 45 seconds at 6.5 m s^-1^. Samples were then centrifuged at 16,000 × *g* for 15 minutes at 4 °C and lysate was transferred to a new tube, followed by a clean-up using Wessel-Flügge precipitation ^88^. The precipitated proteins were dissolved in SDS-PAGE buffer, heated in the water bath for 5 minutes at 95 °C and ran on SDS-PAGE Any-kD Mini-PROTEAN TGX Stain-Free gels (Bio-Rad, California, USA) for 3 minutes. The gel was stained using Coomassie Blue R-250 and visual protein bands were carefully cut into 1×1 mm pieces before proceeding to reduction, alkylation, and digestion into peptides using trypsin. Peptides were concentrated and eluted using C18 stage tips, before dried on a SpeedVac, re-suspended in 0.1% formic acid, and quantified using a Nanodrop One instrument. Finally, the peptide samples were processed using a nano LC-MS/MS coupled to a timTOF Pro mass spectrometer (Bruker, Germany), as described in detail in **Supplementary Text S3**.

#### (meta)genome-centric metaproteomics data analysis

The MS raw data were analysed using FragPipe v16.3, with the search engine MSFragger v3.3^89^ embedded, in addition to Philospopher v4.0.0 ^90^. Here, a protein sequence catalogue retrieved from the 291 recovered MAGs, viral scaffolds (described in **Supplementary Text S4**), and publicly available rumen eukaryote genomes previously described in Andersen, Altshuler ^91^, were used as a sample specific database. Common contaminants such as human keratin, bovine serum albumin and host genome (*Ovis Aries*) were included in the database, as well as decoy sequence entries based on the reverse protein sequences. Carbamidomethylation was used as fixed modification, while oxidation of methionine and protein N-terminal acetylation were added as variable modifications. Trypsin was used as digestive enzyme, with maximum one missed cleavage allowed. For Label-Free-Quantification (LFQ), IonQuant was applied with FDR-controlled Match-between-runs (MBR) enabled ^92^. Protein groups detected from FragPipe were uploaded to Perseus v1.6.15.0 ^93^, for further downstream analyses. First, protein groups identified as contaminants and host proteins were removed, and the intensities were Log2(x) transformed. Furthermore, a protein group needed to be detected in minimum three of eight biological replicates in at least one of the experimental diet groups (control and/or 5% inclusion of *S. latissima*) to be considered. For statistical analysis, missing values were imputed from a 2.8 downshifting from the normal distribution. Protein groups with significant changes in LFQ intensities were identified using a two-sided Student’s T-test (p-value < 0.05).

#### Biochemical analysis of PLs

PL6 enzymes from *Bacteroidota spp.* alginate AULs, and an outgroup OTU, *Brevundimonas Bullata* that does not have a traditional SusCD system, were selected for their taxonomic and phylogenetic diversity within the PL6 family. Selected PL6 genes were trimmed to remove the signal peptide and synthesized into pET28a with a C-terminal His_6_-tag and codon optimized for *E. coli* production (Biobasic). Expression plasmids were transformed into BL21 (DE3) Tuner competent cells (Novagen) and grown to an OD_600nm_ 0.6-0.8 in LB Miller broth containing 50 μg mL^−1^ kanamycin before induction with 1 mM IPTG at 18 °C overnight. Induced cultures were pelleted at 6,500 x *g* for 20 min and lysed using a combination of lysis buffer (20 mM Tris pH 8.0, 500 mM NaCl, 0.1 mg mL^-1^ lysozyme) and sonication (2 min of 1 s intervals of medium intensity sonic pulses at a power setting of 4.5 – Heat systems Ultrasonics Model W-225). Cell lysate was centrifuge at 17,500 x *g* for 1 h and filtrate was purified using Ni-NTA resin columns (Cytiva) and immobilized metal affinity chromatography. Recombinant protein was eluted with an imidazole gradient in 20 mM Tris, pH 8. Recombinant PL6 enzymes were concentrated and dialyzed into 20 mM Tris pH 8.0, 500 mM NaCl, and 2% glycerol using 30 kDa cut-off Amicon Ultra centrifugal concentrators (Millipore Sigma).

50 nM of recombinant PL6 enzymes was incubated with 5 mg mL^-1^ brown seaweed alginate (Sigma) in 20 mM of varying buffers to determine pH optima at 37 °C: citrate-phosphate (pH 4-8.5), CHES (pH 8.5-10), Glycine (pH 8.5-11), and CAPs (pH 10-11). PL6 β-elimination reaction and the generation of C4-C5 unsaturation can be monitored with absorbance at 232 nm. Reactions were run for 10 minutes while continuously monitoring absorbance at 232 nm at 30 s intervals using a SpectraMax ID3 plate reader. Initial velocities of PL6 enzyme activity on brown seaweed alginates were determined using eight concentrations of substrate at the optimal pH for each enzyme. Rates of product formation were calculated in GraphPad Prism (9.0.2) using the extinction coefficient 6,150 M^-1^ cm^-1^ to convert absorbance to product concentration ^94^. Due to substrate complexity in molecular weight and M/G ratio and organization, velocities were unable to be fit to the Michaelis Menten equation.

Reactions were ethanol precipitated by incubating in 50% EtOH overnight at −20 °C before centrifugation at 12,000 x *g* for 10 min and retrieving the supernatant to remove larger, undigested alginate polymers. Samples were dried on a SpeedVac to remove EtOH and resuspended in ultrapure 18 MΩ cm^-1^ H_2_O. Digests were run on thin-layer chromatography using a mobile phase of 2:1:1 butanol: acetic acid: H_2_O, and visualized using an orcinol solution (70:3 acetic acid: sulfuric acid, and 1% orcinol). HPAEC-PAD was performed on a Dionex ICS-3000 using a 3 x 150 mm CarboPac PA20 column (Thermo Scientific). Digests were eluted at a 0.35 mL min^-1^ flowrate at 30 °C in a background of 100 mM NaOH in an increase sodium acetate gradient (0 to 20 min, 0 to 0 M; 20 to 80 min, 0 to 1 M).

Liquid chromatography was performed on a Vanquish ultra-high performance liquid chromatography (UHPLC) system (Thermo Scientific). Alginate oligosaccharide samples were prepared in water and injected in a volume of 10 µL at a concentration of 200 µg/mL. Separation of the digested oligosaccharides was achieved using an Acquity UPLC BEH Amide (HILIC) Column, 130Å, 1.7 µm, 2.1 mm X 150 mm (Waters) at a flow rate of 300 µL/min at 30°C, using a gradient as shown in **Supplementary Table S4** ^95^. Electrospray ionization mass spectrometry (ESI-MS) was performed on an Orbitrap Fusion Tribrid system (Thermo Scientific) in negative ion mode. Mass spectra parameters are shown in **Supplemental Table S5**. To select ions for MS2 experiments, an intensity threshold filter was employed with a minimum intensity of 25,000 and maximum intensity of 1E+20, and a dynamic exclusion filter was used after 1 times for 2.5 s with default mass tolerance values. Higher-energy collisional dissociation (HCD) was employed to generate fragments. MS spectra were analyzed using Xcalibur and Freestyle software packages (Thermo Scientific), and product ions were identified based on both manual interpretation and comparison to previous studies ^40^.

#### Fluorescent Polysaccharides (FLAPS)

A hot water extraction (HWE) was performed on the dried, ball-milled *S. latissima* sample by incubating 1 g of sample in 40 mL distilled water for 8 h at 70 °C. Samples were centrifuged (3,000 × *g*, 10 min) and the supernatant was transferred to a new tube, and the hot water incubation was repeated twice more. Supernatants from the same sample were pooled and freeze-dried. The dried *S. latissima* HWE was de-starched by incubation with α-amylase (E-BLAAM, Megazyme) in 100 mM maleic acid buffer (pH 6.0) containing 100 mM NaCl and 3 mM CaCl_2_ for 8 h at 70 °C, followed by an incubation with amyloglucosidase (E-AMGDPD, Megazyme) for 4 h at 50 °C. Resulting de-starched samples were extensively dialyzed (3,500 Da MWCO) against distilled water, before being freeze-dried. Fluorescently labelled *S. latissima* HWE (FLA-SLAT) was produced using a previously defined protocol ^48^, where the purified FLA-SLAT was freeze-dried, covered in aluminum foil and stably stored at −20 °C until further use. For FLA-SLAT incubations, 1 mL from each RUSITEC vessel was collected at the day 15 time point, and were centrifuged (5,000 × *g*, 10 min). Pellets were suspended in 1 mL phosphate buffered saline (PBS; pH 7.4) and washed twice with PBS before a final resuspension in 2 mL PBS. 50 µL of each resuspended RUSITEC microbial community sample was incubated with 50 µL of 0.4% FLA-SLAT for 1 day within an anaerobic chamber (atmosphere: 85% N_2_, 10% CO_2_, 5% H_2_, at 37 °C), where 40 µL aliquots were taken at the 1 h and 1 day time points. The 1 h and 1 day aliquots were immediately centrifuged (5,000 × *g*, 10 min), and the pellet was fixed with 4% (v/v) formaldehyde overnight at 4 °C. Fixed samples were centrifuged (5,000 × *g*, 10 min) and pellets were washed twice with PBS before being resuspended in 1 mL PBS and stored at 4 °C. Samples were diluted 1:10 and filtered onto a 25 mm, 0.2 µm pore size Isopore™ filter (Sigma, USA) using a gentle vacuum of <200 mbar. Dried filter pieces were counterstained with 4’6-diamidino-2-phenylindole (DAPI) and mounted on a glass microscope slide using a 4:1 mixture of Citiflour™ AFI mountant solution to Vectashield® vibrance antifade mounting medium (Vector Laboratories, USA). The fluorescence images were taken using an Echo Revolve R4K (Upright & Inverted Capability) microscope equipped with motorized LED fluorescence light, 5 MP CMOS monochrome camera, a Plan X Apo oil; 1,42 NA, Revolve, 60x oil immersion objective, with LED light cubes DAPI (EX: 385/30 EM: 450/50 DM: 425) and LED light cubes FITC (EX: 470/40 EM: 525/50 DM: 495). SR-SIM images for RUSITEC samples were visualized on a Zeiss ELYRA PS.1 (Carl Zeiss) using 561 and 488m lasers and BP 573-613, BP 502-538 and BP 420-480 + LP 750 optical filters. Z-stack images were taken with a Plan-Apochromat 63 Å∼ /1.4 Oil objective and processed with the software ZEN2011 (Carl Zeiss).

Images were exported to ACMEtool software (M. Seder, Technology GmbH, http://www.technobiology.ch and Max Planck Institute for Marine Microbiology, Bremen), where signals were evaluated according to Bennke, Reintjes ^96^. Cell enumeration counts were plotted in GraphPad Prism (8.0.2) and displayed as relative abundance values.

## Acknowledgements

We thank Cascadia Seaweed Corporation for the generous donation of *S. latissima*. Research at AAFC was supported through an AgriScience Project (ASP-207; J-002817) and the Prize for Outstanding Achievement in Science (AAFC) awarded to D.W.A. (J-003135). LC-ESI-MS data was collected at the University of Lethbridge Magnetic Resonance Facility with support from Tony Montina, Vincent Weiler, Maurice Needham, and Carl Holland. L.H.H. and A.F. are grateful for support from The Research Council of Norway (project no: 302639). P.B.B. is grateful for support from the Novo Nordisk Foundation (0054575—SuPAcow) and the Australian Research Council (Future Fellowship: FT230100560). The lamb feeding experiment was financed by The Research Council of Norway; BIOTEK2021/Havbruk Biofeed (project no. 229003) and Foods of Norway a Centre for Research Based Innovation (project no. 237841). The sequencing service of rumen samples was provided by the Norwegian Sequencing Centre (www.sequencing.uio.no), a national technology platform hosted by the University of Oslo and supported by the ‘Functional Genomics’ and ‘Infrastructure’ programs of the Research Council of Norway and the Southeastern Regional Health Authorities. The authors acknowledge the Orion High Performance Computing Center at the Norwegian University of Life Sciences and Sigma2—the National Infrastructure for High Performance Computing and Data Storage in Norway for providing computational resources that have contributed to meta-omics computations reported in this paper. T.R.P. was supported by the Canada Research Chair program. Mass spectrometry-based proteomic analyses were performed by The MS and Proteomics Core Facility, Norwegian University of Life Sciences (NMBU). This facility is a member of the National Network of Advanced Proteomics Infrastructure (NAPI), which is funded by the Research Council of Norway INFRASTRUKTUR-program (project number: 295910). We are thankful for the generous access provided by the Molecular Ecology Department of the Max Planck Institute for Marine Microbiology for microscopy. G.R. was supported through funding from the Deutsche Forschungsgemeinschaft (DFG, German Research Foundation; project number 496343779).

## Contributions

A.F., J.P.T., D.W.A. and L.H.H. conceived the study, performed the primary analysis of the data, and generated figures. D.W.A., T.R.P., and L.H.H. supervised the work. S.A.T. and D.W.A. designed and performed the RUSITEC analysis. P.B.P. and L.H.H. proposed and performed the lamb omics analysis. L.T.M., M.Ø., and A.K. designed, performed and/or analyzed the lamb feeding trial. T.J. and A.L. extracted lamb rumen DNA. J.P.T., and M.L.K. contributed to the RUSITEC data analysis. B.B. and X.X. contributed to AIR preparation and linkage analysis for *S. latissima*. G.R., L.K., and M.L.K. generated and analyzed FLAPS results. J.P.T. and A.Y.S. characterized alginate lyases, K.E.L. generated and analyzed LC-MS. A.F., J.P.T., D.W.A., and L.H.H. wrote the paper, with valuable inputs and edits from P.B.P. and G.R. All authors reviewed and approved the final manuscript.

