## Supplementary information for "Microbial alginate foraging is conserved in geographically and taxonomically distinct ruminant microbiomes"

**CONTENT:**

I. Supplementary Figures
II. Supplementary Tables
III. Supplementary Text

Metagenome-Assembled Genome (MAG) information and metaproteomics data are provided in Extended Data 1 and Extended Data 2, respectively.

#### Supplementary Figures

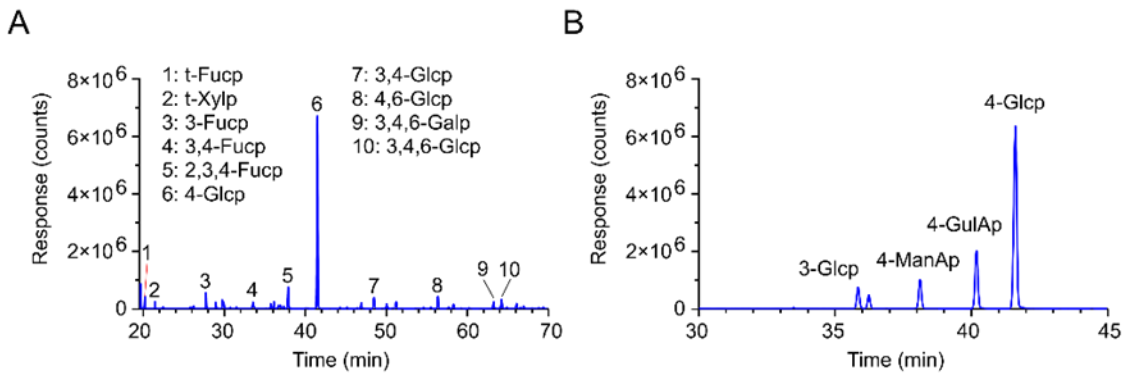

**Supplementary Figure S1: Linkage analysis chromatographs of *S. latissima***. Total ion current chromatograms of *S. latissima* with (A) standard methanolysis and (B) carboxyl reduced methanolysis with major PMMAs labelled.

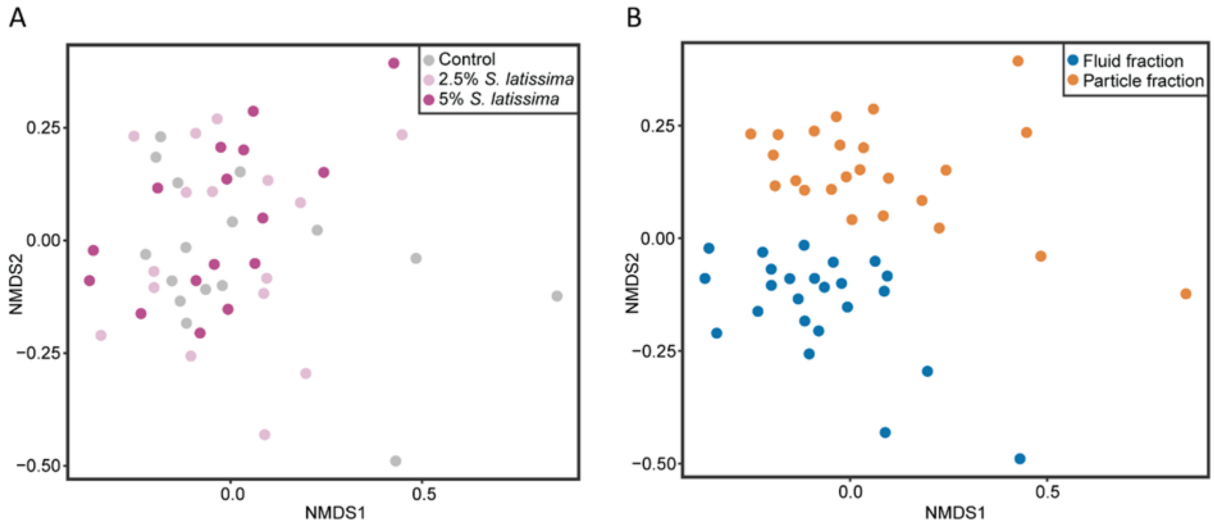

**Supplementary Figure S2: *In vivo* community analysis of lamb rumen microbial communities (16S rRNA gene) from dietary groups supplemented with 2.5% and 5% *S. latissima.*** (A) Non-metric multi-dimensional scaling (NMDS) plot comparing the community composition in samples taken from control (grey), 2.5% (light purple) and 5% (dark purple) *S. latissima* dietary groups. B) The NMDS plot colored by sample fraction, either fluid (blue) or particle (orange).

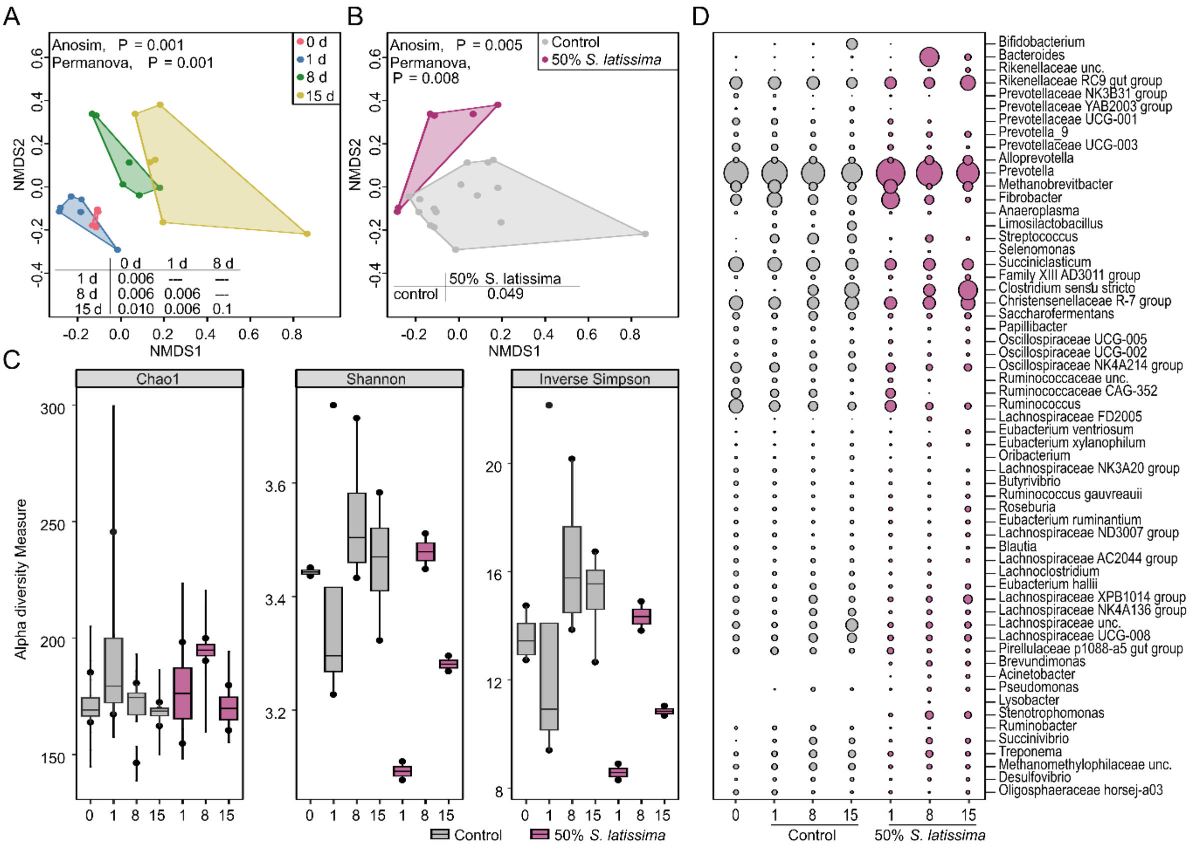

**Supplementary Figure S3. *Ex vivo* community analysis of RUSITEC rumen microbial communities (16S rRNA gene) supplemented with 50% *S. latissima*.** A) Non-metric multi-dimensional scaling (NMDS) plot comparing the community composition at different time points: 0 d (pink), 1 d (blue), 8 d (green), and 15 d (yellow). B) NMDS plot comparing the effect each treatment has on the community composition where control (grey) and 50% *S. latissima* (purple). C) Alpha diversity (Chao1, Shannon, and inverse Simpson indices) in RUSITEC rumen microbial communities supplemented with 50% *S. latissima* over time. D) Bubble plot of different bacterial and archaeal genera that reached a minimum relative read abundance of 1% in control (grey) and 50% *S. latissima* (purple) RUSITEC rumen microbial communities.

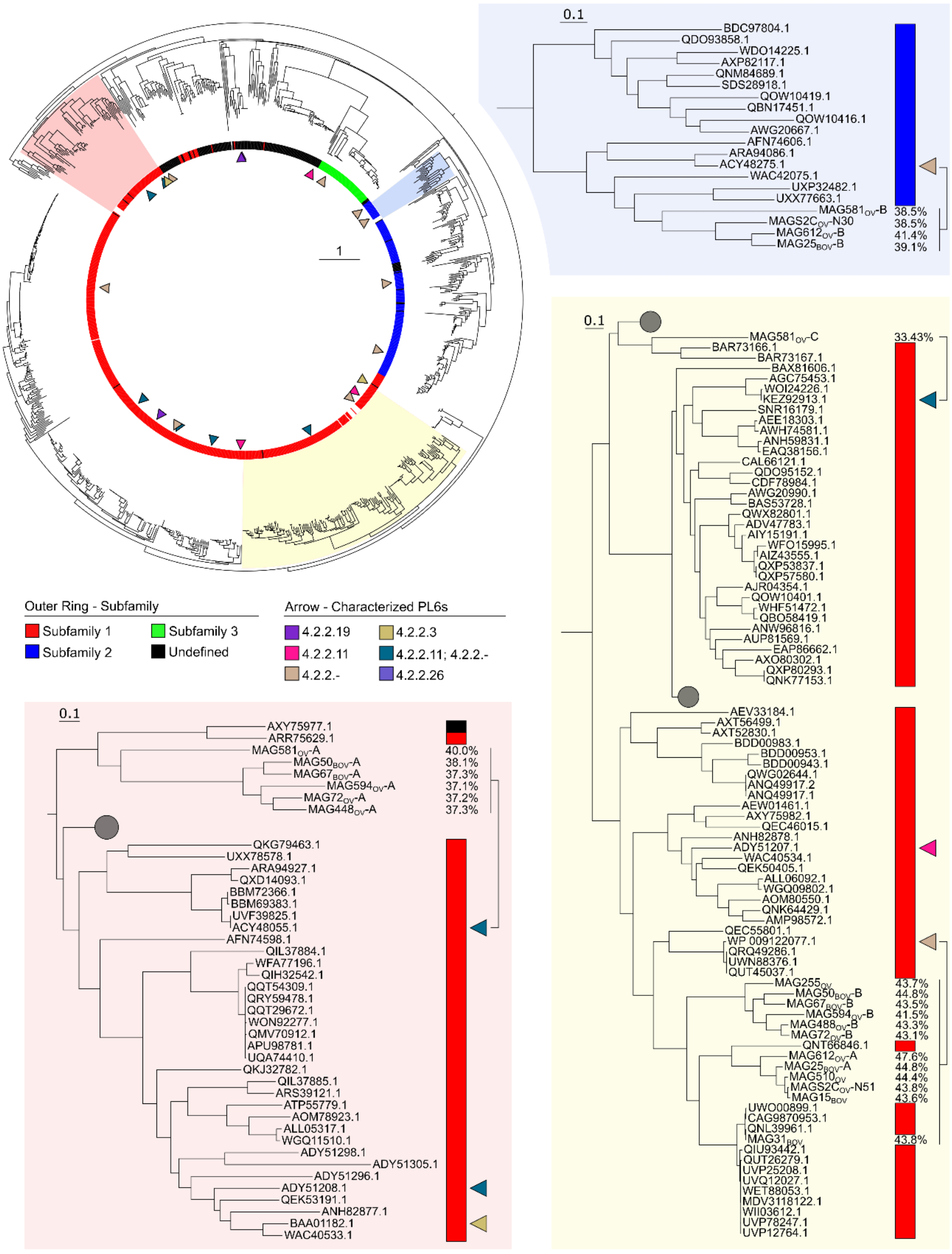

**Supplementary Figure S4: Homology of *Bacteroidota* spp. PL6 members within this study.** SACCHARIS phylogeny of PL6 members within CAZy and *Bacteroidota* spp.. The full phylogeny (top left) highlights the three clades *Bacteroidota* spp. PL6s within this study was found (red – clade 1; blue – clade 2; yellow – clade 3). Clades where further pruned to identify closest related PL6 members (color highlighted as above), and the PL6 members within each clade was compared to the closest related characterized PL6 member. Clade 1 – ACY48055.1 (*Rhodothermus marinus* DSM 4252), clade 2 – ACY48275.1 (*Rhodothermus marinus* DSM 4252), and clade 3 - KEZ92913.1 (*Nonlabens ulvanivorans* PLR) and WP_009122077.1 (*Bacteroides clarus* YIT 12056).

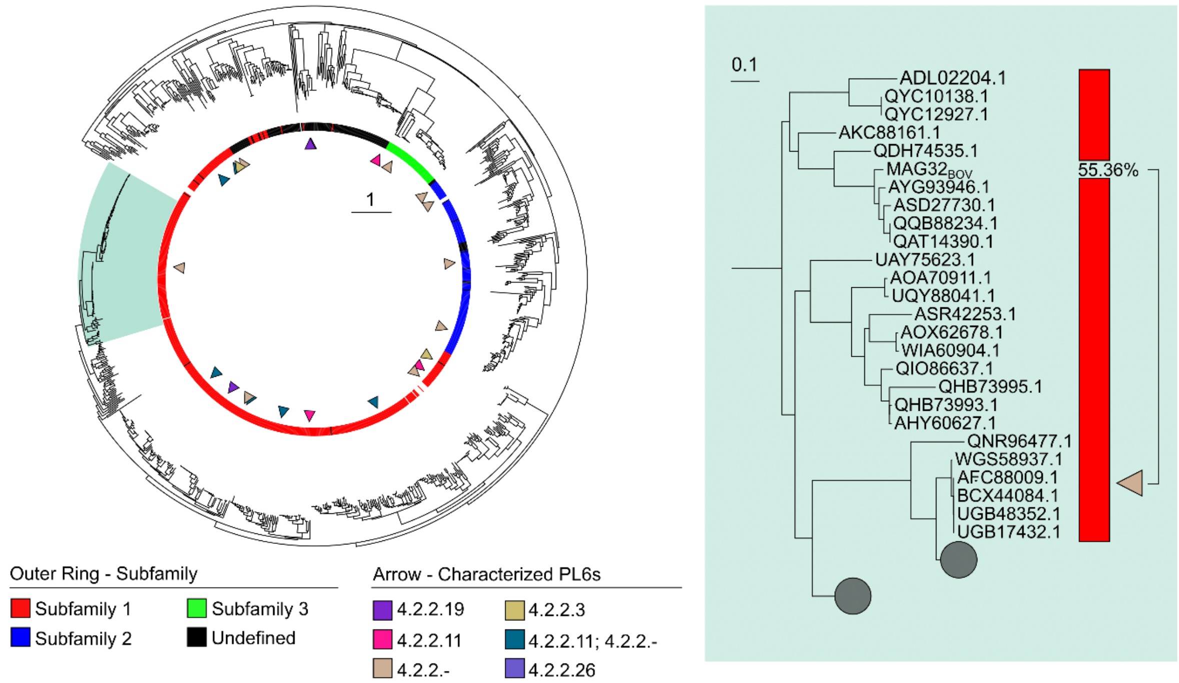

**Supplementary Figure S5: Homology of *B. bullata* PL6 towards other PL6 members.** SACCHARIS phylogeny of PL6 members within CAZy and $MAG32_{BOV}$ PL6 member. The full phylogeny (left) highlights the $MAG32_{BOV}$ clade, which is further pruned (right) to display closest related PL6 members. The identity between $MAG32_{BOV}$ PL6 and the closest characterized PL6 member (AFC88009.1 - *Stenotrophomonas maltophilia* KJ-2) is displayed.

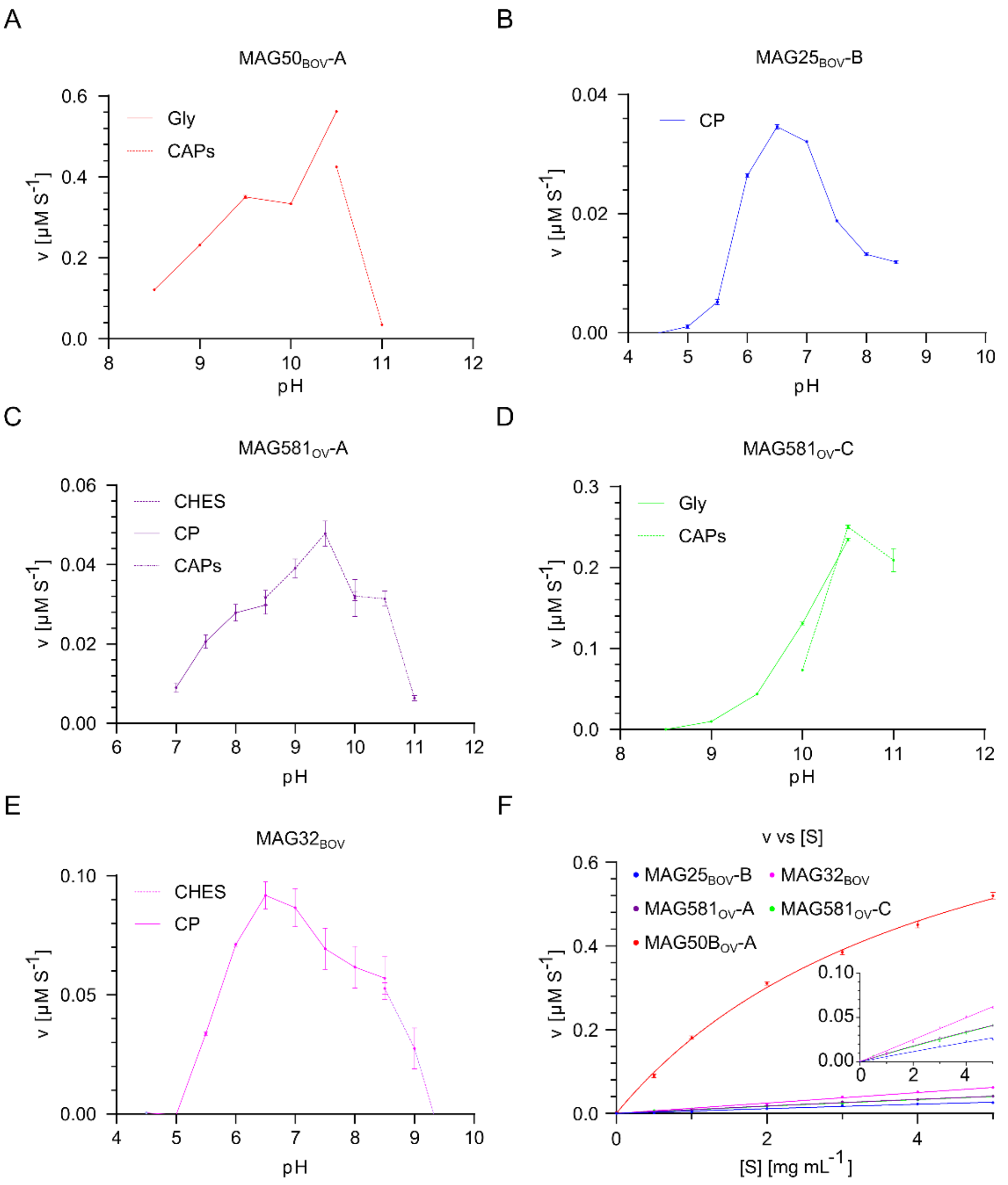

**Supplementary Figure S6: pH and initial velocities of PL6 members on brown seaweed alginate.** A-E) pH optima of PL6 within this study. Optima was determined using product formed (µM – calculated from absorbance at 232 nm) per second for 10 minute reactions. Buffers: Gly – Glycine, CAPs - 3-(Cyclohexylamino)propane-1-sulfonic acid, CP – Citrate-phosphate, CHES - N-cyclohexyl-2-aminoethanesulfonic acid. F) initial velocities of PL6s within study. Velocities were determined using product formed (µM – calculated from absorbance at 232 nm) per second against substrate concentration (mg $mL^{-1}$) for 10 minute reactions.

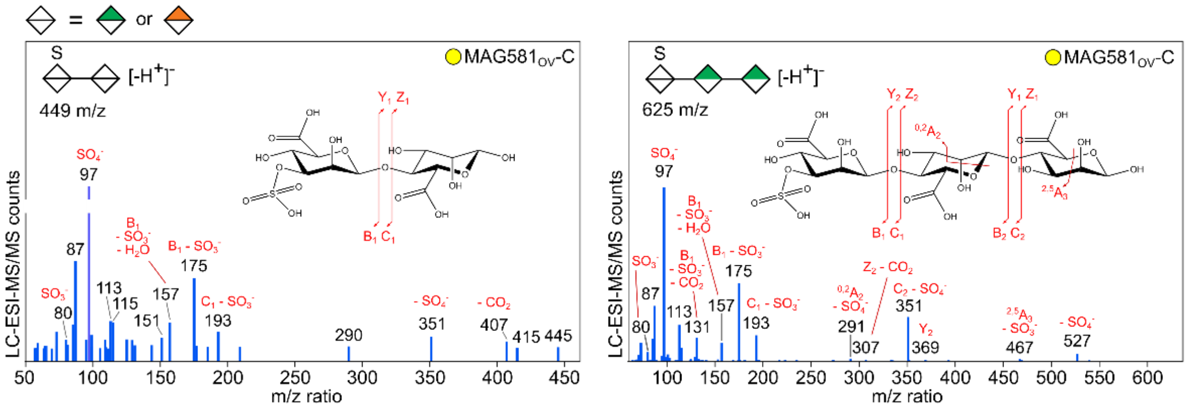

**Supplementary Figure S7: Sulfated alginate oligosaccharide enzyme products as identified by LC-ESI-MS/MS.** Commercial alginic acid was incubated with MAG581-C enzyme and analyzed by LC-ESI-MS/MS. Oligosaccharide species for ions of *m/z* 449 and 625 were identified based on MS2 fragmentation data. Monosaccharide symbols are displayed according to the Symbol Nomenclature for Glycans system [1]. For monosaccharides unable to be discerned by LC-MS between mannuronate and guluronate, white symbols are used.

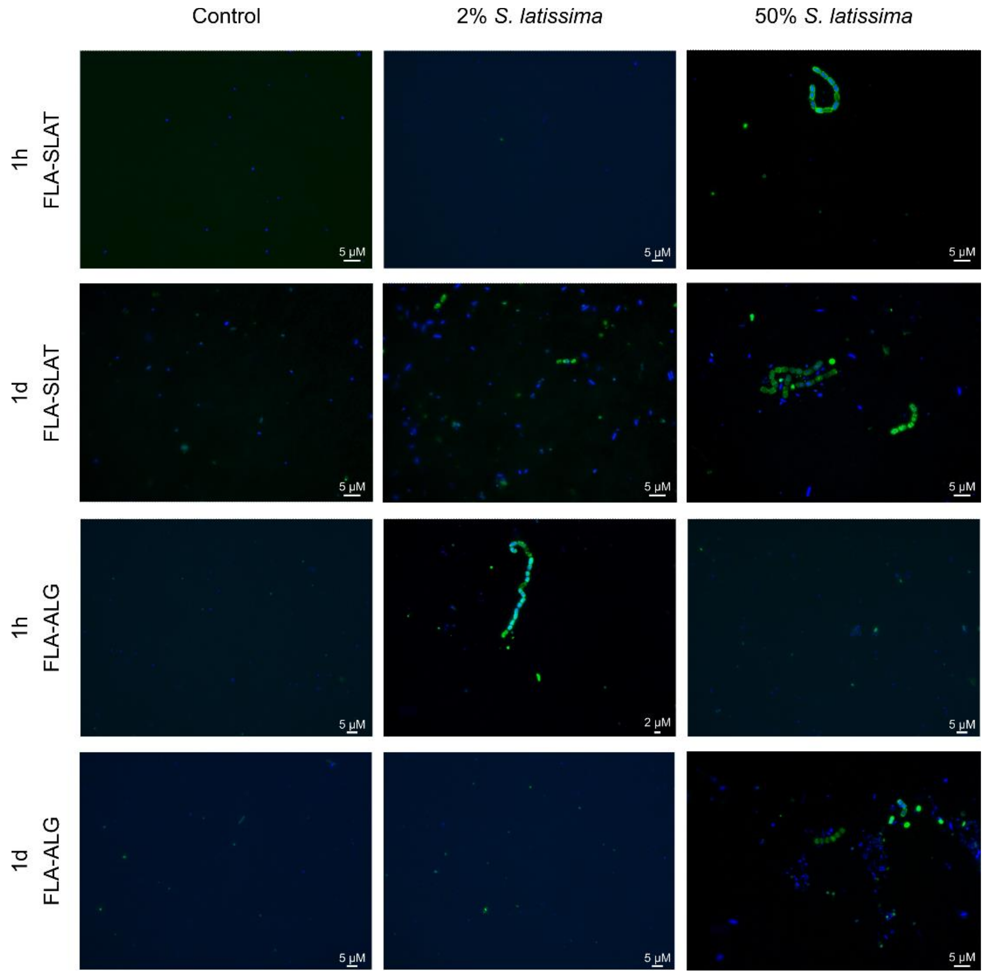

**Supplementary Figure S8: Uptake of fluorescently labelled polysaccharides by RUSITEC communities.** RUSITECs containing 0% (control), 2%, or 50% *S. latissima* were sampled on day 15 and incubated with fluorescently labelled alginate (FLA-ALG) or *S. latissima* extract (FLA-SLAT) for 1h or 1d. Samples were co-stained with DAPI and imaged by an Echo Revolve R4K microscope.

### Supplementary Tables

**Supplementary Table S1:** Relative abundances (Mol%) of glycosidic linkages identified from the cell walls of *Saccharina latissima* via GC-MS; n=3.

| PMAA | Avg. Mol % | PMAA | Avg. Mol % | PMAA | Avg. Mol % |
|---|---|---|---|---|---|
| 3-Araf | Trace | 4-Glcp | 35.0±6.4 | t-Rhap | Trace |
| 5-Araf | Trace | 6-Glcp | Trace | 2-Rhap | Trace |
| t-Fucp | 2.1±0.2 | 2,3-Glcp | Trace | 3-Rhap | Trace |
| 2-Fucp | 1.2±0.1 | 3,4-Glcp | 2.7±0.2 | 4-Rhap | Trace |
| 3-Fucp | 3.2±0.2 | 3,6-Glcp | Trace | 2,3-Rhap | Trace |
| 4-Fucp | 1.7±0.4 | 4,6-Glcp | 2.5±0.4 | 2,4-Rhap | Trace |
| 2,3-Fucp | 1.1±0.2 | 2,3,6-Glcp | Trace | 3,4-Rhap | Trace |
| 2,4-Fucp | 0.8±0.1 | 2,4,6-Glcp | Trace | 2,3,4-Rhap | Trace |
| 3,4-Fucp | 2.0±0.6 | 3,4,6-Glcp | 2.6±0.8 | t-Xylp | 1.5±0.3 |
| 2,3,4-Fucp | 5.1±1.0 | 2,3,4,6-Glcp | 0.8±0.3 | 2-Xylp | Trace |
| t-Galp | Trace | t-Manp | Trace | 3-Xylp | Trace |
| 2-Galp | Trace | 2-Manp | 1.2±0.2 | 4-Xylp | Trace |
| 4-Galp | Trace | 3-Manp | Trace | 2,4-Xylp | Trace |
| 6-Galp | Trace | 4-Manp | Trace | 3,4-Xylp | Trace |
| 3,4-Galp | 1.4±0.2 | 2,3-Manp | Trace | 2,3,4-Xylp | Trace |
| 3,6-Galp | 1.1±0.2 | 2,4-Manp | 1.2±0.2 | t-GalpA | Trace |
| 4,6-Galp | Trace | 2,6,Manp | Trace | 2,4-GlcpA+2,4-GalpA | Trace |
| 2,3,6-Galp | Trace | 3,4-Manp | Trace | t-GlcpA | Trace |
| 2,4,6-Galp | Trace | 3,6-Manp | Trace | 3-GlcpA | 2.6±0.1 |
| 3,4,6-Galp | 1.8±0.4 | 4,6-Manp | Trace | 4-GlcpA | 2.4±0.1 |
| 2,3,4,6-Galp | Trace | 2,3,6-Manp | 0.7±0.2 | t-GulpA | Trace |
| 2,4-Glcp+2,4-Galp | 1.6±0.3 | 2,4,6-Manp | Trace | 4-GulpA | 10.1±1.8 |
| t-Glcp | Trace | 3,4,6-Manp | Trace | t-ManpA | Trace |
| 3-Glcp | 0.9±0.0 | 2,3,4,6-Manp | Trace | 4-ManpA | 4.6±0.6 |

**Supplementary Table S2.** The effect of 2.5% and 5% *S. latissima* on dry matter intake and apparent total tract digestibility of nutrients with Norwegian White lambs

| | Control | 2.5% *S. latissima* | 5% *S. latissima* | SEM | P-value |
|---|---|---|---|---|---|
| DMI, kg/d | 1.55 | 1.53 | 1.55 | 0.04 | 0.66 |
| *Digestibility (ATTD), %* | | | | | |
| DM | 81.1 | 80.0 | 78.9 | 2.63 | 0.27 |
| OM | 81.7 | 80.5 | 79.4 | 2.66 | 0.19 |
| CP | 83.5 | 81.6 | 79.6 | 2.57 | 0.012 |
| NDF | 78.5 | 77.7 | 76.2 | 3.13 | 0.36 |
| Starch | 99.3 | 99.1 | 99.0 | 0.28 | 0.13 |
| Ash | 74.4 | 74.2 | 74.0 | 2.56 | 0.96 |

DMI, Dry matter intake; ATTD, apparent total tract digestibility of nutrients, DM, dry matter; OM, organic matter; CP, crude protein; NDF, neutral detergent fiber.

**Supplementary Table S3:** The effect of 2% *S. latissima* on the degradability of nutrients and total volatile fatty acid production in the RUSITEC system

| | Control | 2% *S. latissima* | SEM | P-value |
|---|---|---|---|---|
| *Degradability, %* | | | | |
| DM | 57.1 | 58.4 | 0.80 | <0.001 |
| OM | 58.0 | 57.3 | 1.67 | 0.45 |
| CP | 69.3 | 69.0 | 0.85 | 0.77 |
| NDF | 36.1 | 35.9 | 2.75 | 0.87 |
| Total VFA, mmol | 68.4 | 68.3 | 1.82 | 0.95 |

DM, dry matter; OM, organic matter; CP, crude protein; NDF, neutral detergent fiber; VFA, volatile fatty acids

**Supplementary Table S4:** LC-MS Gradient conditions for separation of alginate oligosaccharides.

| Time (min) | A (%)<br>10 mM ammonium formate<br>50 mM formic acid<br>80% acetonitrile<br>20% water | B (%)<br>10 mM ammonium formate<br>50 mM formic acid<br>20% acetonitrile<br>80% water |
|---|---|---|
| 0 | 100 | 0 |
| 1 | 70 | 30 |
| 60 | 20 | 80 |
| 61 | 20 | 80 |
| 67 | 100 | 0 |
| 68 | 100 | 0 |
| 75 | 100 | 0 |

**Supplementary Table S5:** Parameters for ESI-MSn on the Orbitrap Fusion Tribrid.

| Parameter (units) | Value |
|---|---|
| ***ESI*** | |
| Spray voltage: negative ion (V) | 2500 |
| Sheath Gas (Arb) | 35 |
| Aux Gas (Arb) | 10 |
| Sweep Gas (Arb) | 1 |
| Ion Transfer Tube Temp (°C) | 325 |
| Vaporizer Temp (°C) | 250 |
| **MS** | |
| Detector Type | Orbitrap |
| Orbitrap Resolution | 120K |
| Mass Range | Normal |
| Scan Range (m/z) | 150-2000 |
| RF Lens (%) | 50 |
| **MS2** | |
| Collision Energy Type | Normalized |
| Isolation Mode | Quadrupole |
| Activation Type | HCD |
| Collision Energy Mode | Stepped |
| Collision Energies (%) | 30,45,60,80 |
| Detector Type | Orbitrap |
| Orbitrap Resolution | 30K |

### Supplementary Text

##### Text S1: Digestibility and degradability impacts of *S. latissima* in diets

Evaluating the digestibility of seaweed in animal diet is crucial for understanding its effect on the overall ruminal digestive activity and nutrient utilization. Thus, ruminal kelp digestion was assessed both in the *in vivo* lamb feeding and the *in vitro* bovine RUSITEC experiments.

For the lamb experiment, mean dry matter intake (DMI, kg/d) was not affected by the inclusion of *S. latissima* to diets. However, *S. latissima* inclusion at 5% (DM, basis) reduced apparent total tract digestibility of crude protein by 4.67% relative to the control diet (**Supplementary Table S1**). Digestibility of other tested dietary components were not significantly affected despite numerical differences. For the RUSITEC experiment, inclusion of 2% *S. latissima* increased the *in vitro* degradability of DM by 2.24%. There were no effects of *S. latissima* inclusion on organic matter, crude protein, neutral detergent fiber, or production of total volatile fatty acids (**Supplementary Table S2**).

##### Text S2: 16S rRNA gene sequencing

Taxonomic assigned ASV from the lamb rumen microbiome were combined with variant abundance table and processed using Phyloseq v1.46.0 [2]. Reads flagged as Eukaryota, Chloroplast or Mitochondria were removed before the sequences were normalized for downstream analysis. The beta diversity was investigated using a Non-metric Multidimensional scaling (NMDS) analysis based on Bray-Curtis dissimilarity in vegan [3].
RUSITEC community analysis, statistics, and plotting of the Bracken output files were performed using R v4.2.2 in R-studio v2022.02.3 [4] with the packages: phyloseq [2], ggplot2 [5], picante [6], rioja [7] and vegan [3]. Each sample was rarefied to 28,530 random reads using the "rarefy_even_depth(sample.size = 0.9*raremax)" function of phyloseq [2] prior to alpha- and beta-diversity analyses. NMDS analyses based on Bray-Curtis dissimilarity indices were performed using the "vegdist()" function of vegan [3] to visualize the separation of RUSITEC microbial communities between the different treatments and time points. Stress was determined to be 0.097 with the "metaMDS()" function [3], and a Shepard diagram was used to determine the non-metric ($R2= 0.991$) and linear fit ($R2= 0.968$) between the observed dissimilarity. Statistical evaluation of the RUSITEC microbial communities was performed with an analysis of similarity (ANOSIM) using distance matrices ["anosim()" function, distance =

bray, 999 permutations] and a permutational multivariate analysis of variance (PERMANOVA) using distance matrices [“vegdist()” function, method = bray, 999 permutations] [3]. This analysis was followed by a pairwise comparison of the treatment or time point effect [pairwise.perm.manova()” function, method = Euclidian, “999 permutations] of RVAideMemoire [8]. Observed OTUs, Shannon, and inverse Simpson indices were calculated using the “estimate_richness()” function of phyloseq , as a measure of microbial community alpha-diversity. Changes in alpha-diversity were assessed in GraphPad Prism v8.0.2 using the Kruskal-Wallis test with a post-hoc Dunn’s multiple comparisons.

**Text S3: Metaproteomics**

After protein extraction, peptide samples were processed using a nano LC-MS/MS coupled to a timTOF Pro mass spectrometer (Bruker, Germany). The peptides were separated by an Aurora C18 reverse-phase (1.6 µm, 120 Å) 25 cm x 75 µm analytical column with an integrated emitter (IonOpticks, Melbourne, Australia). The temperature of the column was kept at 50 °C using the integrated oven. Equilibration of the column was performed before the samples were loaded (equilibration pressure 800 bar). The flow rate was set to 300 nl $min^{-1}$ and the samples were separated using a solvent gradient from 2% to 25% solvent B over 70 minutes, and to 37% over 9 minutes. The solvent composition was then increased to 95% solvent B over 10 min and maintained at that level for an additional 10 min. In total, a run time of 99 min min was used for the separation of the peptides. Solvent A is 0.1% (v/v) formic acid in milliQ water, while solvent B is 0.1% (v/v) formic acid in LC-MS grade acetonitrile. The timsTOF Pro was run in positive ion data dependent acquisition PASEF mode with the control software Compass Hystar v5.1.8.1 and timsControl v1.1.19 68. The acquisition mass range was set to 100 – 1700 m $z^{-1}$. The TIMS settings were: 1 $K0^{-1}$ Start 0.85 V·s $cm^{-2}$ and 1 $K0^{-1}$ End 1.4 V·s $cm^{-2}$, ramp time 100 ms, ramp rate 9.42 Hz, and duty cycle 100%. The capillary voltage was set at 1400 V, dry gas at 3.0 l $min^{-1}$, and dry temp at 180 °C. The MS/MS settings were the following: number of PASEF ramps 10, total cycle time 0.53 sec, charge range 0-5, scheduling target intensity 20,000, intensity threshold 2,500, active exclusion release after 0.4 min, and CID collision energy ranging from 27-45 eV.

**Text S4: Recovery of virome scaffolds from lamb rumen**

In addition to the prokaryote population, we attempted to recover viral content from the lamb metagenome co-assembly. This was done following the viral sequence identification

SOP v3 (DOI:dx.doi.org/10.17504/protocols.io.bwm5pc86), with VirSorter2 v2.2.3 [9] and CheckV v0.8.1 [10]. DRAM-v.py was ran to annotate the sequences identified as viral, followed by manual curation according to the SOP. Annotated virome scaffolds are available in FigShare via doi:10.6084/m9.figshare.28024343. As the virome scaffolds were included in the sequence database for metaproteomics analysis, those with protein detection are provided in Extended Data 2. Further exploration of the rumen virome is beyond the scope of this study.
